# A physics-informed hybrid deep learning model for spatiotemporal rice disease prediction using multi-source data

**DOI:** 10.64898/2026.09.03.749105

**Authors:** Zichao Jin, Jiaoru Wang, Wenjiang Huang, Jingcheng Zhang, Huiqin Ma, Roberto Salguero-Gómez

**Affiliations:** College of Artificial Intelligence, Hangzhou Dianzi University, Hangzhou 310018, China; Department of Biology, University of Oxford, Oxford, UK; State Key Laboratory of Remote Sensing and Digital Earth, Aerospace Information Research Institute, Chinese Academy of Sciences, Beijing 100101, China; Food and Agriculture Organization of the United Nations, Rome, Italy

**Keywords:** hybrid modelling, multi-source data fusion, physics-informed deep learning, *Rhizoctonia solani*, rice sheath blight, spatiotemporal forecasting

## Abstract

Accurate, reliable, large-scale disease predictions are essential to ensure rice production. Existing disease prediction models often face a trade-off between interpretability and predictive capability, necessitating the integration of mechanistic knowledge and data-driven learning within a modelling framework. Accordingly, we propose a physics-informed hybrid gated recurrent unit (PI-HGRU) model for spatiotemporal dynamic prediction of rice sheath blight disease, caused by a fungus. Our model embeds differential equations describing disease transmission dynamics into a hybrid gated recurrent unit (HGRU) framework as mechanistic constraints, thereby enabling collaborative modelling between epidemiological processes and data-driven learning. We conducted model training and evaluation using a long-term, multi-source dataset spanning 17 years (2000–2016) and covering 16 major rice-producing provinces in southern China. These rich data include spatiotemporally aligned field disease observations, remote sensing data, meteorological data, and soil property data. In addition, to address the challenges of irregular sampling intervals and inconsistent sequence lengths in disease survey data, we adopted a sliding time-window-based prediction framework. We further conducted a time-window sensitivity analysis to determine appropriate configurations of the input time window and lag time, enabling the model to represent the cumulative and delayed effects of environmental factors. Our PI-HGRU framework substantially outperforms the purely data-driven HGRU baseline model, improving the squared Pearson correlation coefficient (*r*²) by 22.8% while reducing the root mean square error (RMSE) and mean absolute error (MAE) by 10.2% and 18.0%, respectively. Furthermore, analysis of the model’s intermediate variables showed that the transmission rate *β*(*t*) exhibited interpretable relationships with environmental conditions within the input time window, providing a process-related link between environmental drivers and modeled disease transmission dynamics. Overall, our work demonstrates that integrating epidemiological mechanisms into deep learning models in a physics-informed manner can improve predictive accuracy and stability while enhancing model interpretability, highlighting its potential for large-scale disease forecasting and precision disease management.

## 1. Introduction

Rice (*Oryza sativa* L.) is one of the world’s most important staple crops, providing the primary food source for more than half of the global population. Global rice production reached approximately 820 million tonnes in 2024 (FAO, 2025), highlighting its fundamental role in global food security (Yu et al., 2026). However, rice production is frequently threatened by various diseases, many of which are strongly influenced by environmental conditions and exhibit complex spatiotemporal transmission patterns (Ramalingam et al., 2020; Chen et al., 2023). Among these diseases, rice sheath blight (ShB), caused by the fungus *Rhizoctonia solani* Kühn, is one of the most widespread and destructive rice diseases worldwide (Yellareddygari et al., 2014; Shen et al., 2023). In China, ShB occurs extensively in major rice-growing regions, particularly in the Yangtze River Basin and southern rice-producing areas (Li et al., 2021). Due to its wide geographic distribution and strong environmental dependence, ShB exhibits pronounced spatial and temporal heterogeneity, which greatly increases the difficulty of regional-scale disease forecasting. Therefore, developing models capable of effectively forecasting the spatiotemporal dynamics of ShB at regional scales is of significant importance for disease early warning and management (Jensen et al., 2025).

Crop disease forecasting models have rapidly evolved alongside advances in data acquisition technologies and modelling methodologies (Yuan et al., 2023) (Table 1). Early studies primarily relied on rule-based approaches derived from expert knowledge and environmental thresholds (Jensen et al., 2025). In these approaches, disease risk was assessed through logical inference or fuzzy reasoning using key environmental and phenological factors such as temperature, humidity, and crop growth stage (Litschmann et al., 2020). For example, Gonzalez-Dominguez et al. (2016) developed a fuzzy control system for grape downy mildew forecasting, integrating grape phenology, initial infection risk, and secondary infection conditions through IF–THEN expert rules, thereby reproducing expert reasoning with high consistency. Similarly, Singh et al. (2016) proposed a weather-driven forecasting model for potato late blight based on cumulative temperature and humidity thresholds. Across different regions and years, the model achieved a mean absolute error of only 10.48 days and an overall prediction accuracy of 73.6%. Building upon these approaches, statistical modelling methods were subsequently introduced to quantitatively characterize the relationships between environmental variables and disease occurrence (De Wolf et al., 2003; Zhang et al., 2014; Ahmed et al., 2016). For instance, Ahmed et al. (2016) established a stepwise regression model linking potato late blight severity with multiple epidemiological factors. De Wolf et al. (2003) developed a logistic regression model for predicting Fusarium head blight risk using meteorological conditions during critical pre- and post-flowering periods of wheat, achieving prediction accuracies ranging from 62% to 85%.

**Table 1.**
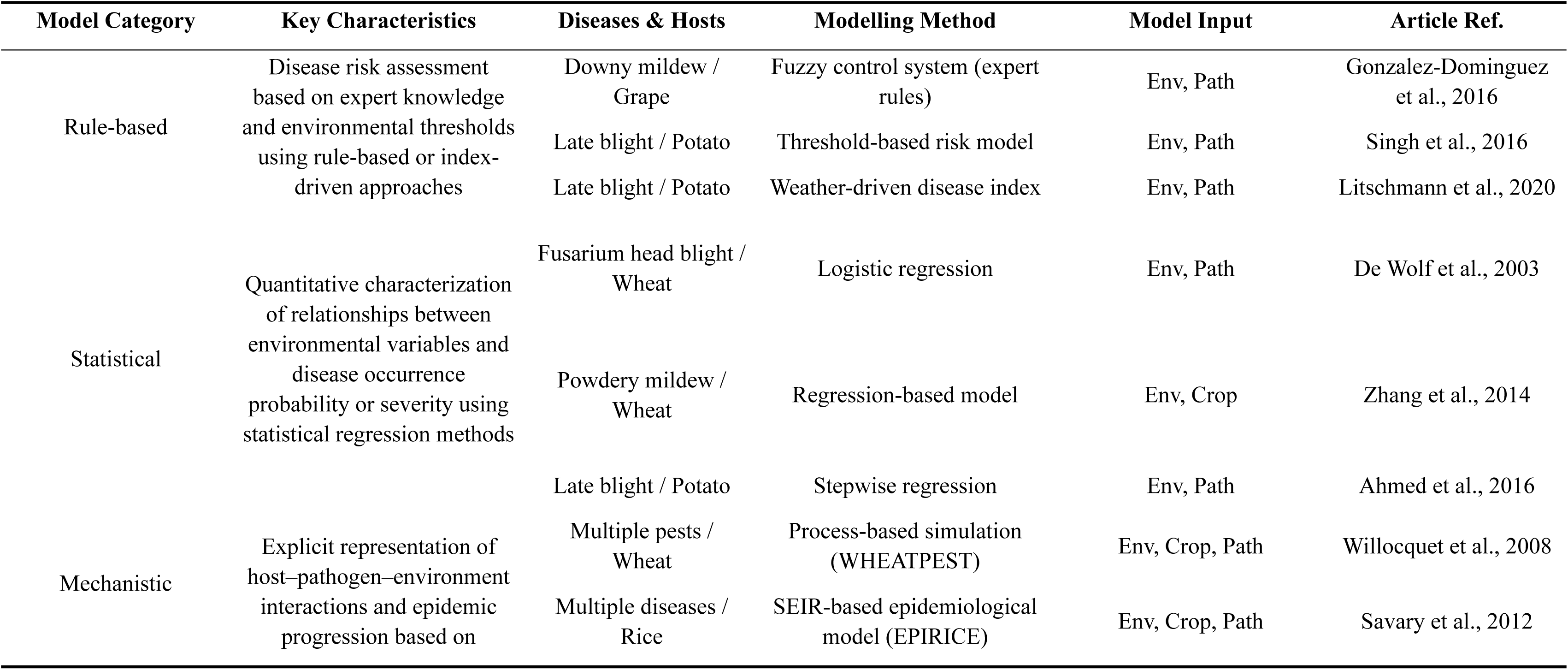
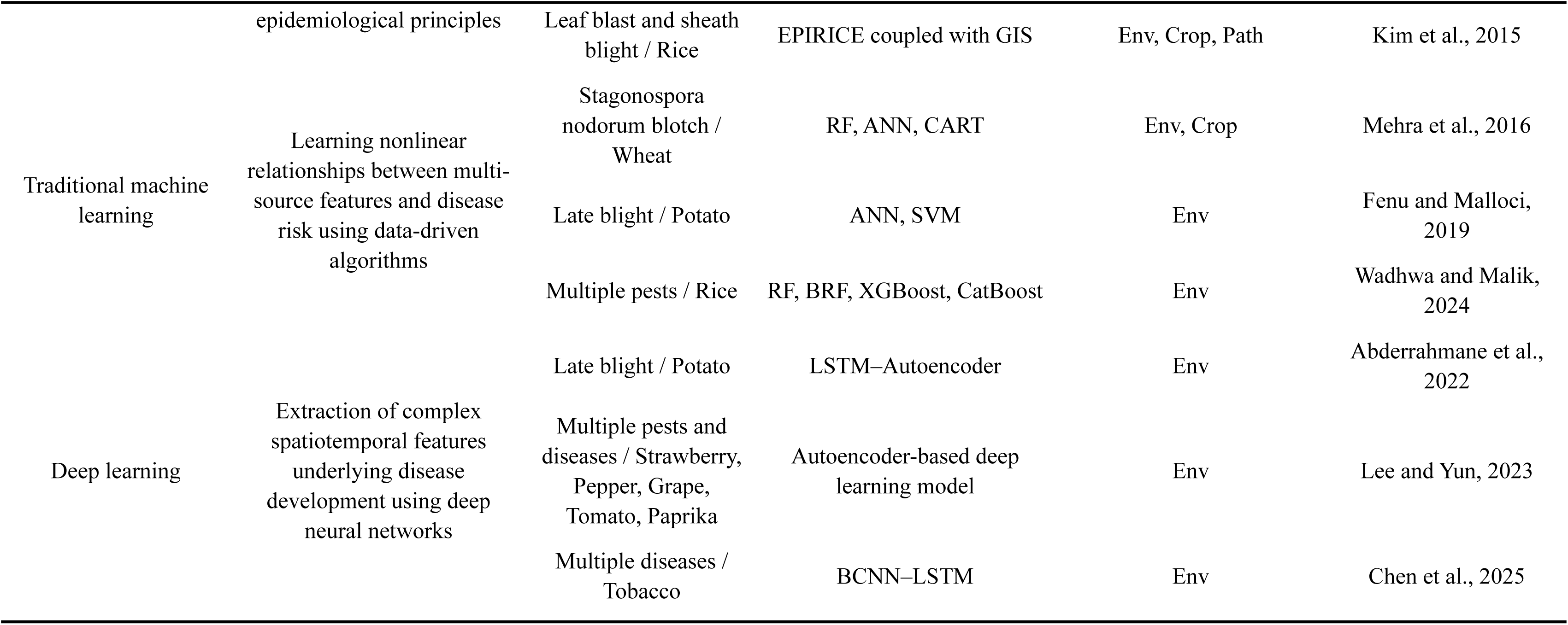
Representative studies illustrating the development of crop disease forecasting approaches, grouped into rule-based, statistical, mechanistic, traditional machine learning, and deep learning approaches. Env denotes environmental data; Crop denotes host/crop physiological or management data; Path denotes pathogen or disease observation data.

Through mathematical representations of disease transmission processes, mechanistic models based on biological and epidemiological principles have also been developed to dynamically simulate disease progression (Willocquet et al., 2008; Savary et al., 2012; Kim et al., 2015). Among these, the Susceptible–Exposed–Infectious–Removed (SEIR) framework has been widely adopted as a classical epidemiological model for delineating infectious disease dynamics in crop disease systems. For example, Savary et al. (2012) developed the EPIRICE model, a generic SEIR-based epidemiological model that successfully simulated the epidemic dynamics of five major rice diseases, including leaf blast, brown spot, bacterial blight, ShB, and tungro. Their simulated disease progression curves showed strong agreement with observed disease progression trajectories, with *R*^2^ values ranging from 0.78 to 0.99. Building on this framework, Kim et al. (2015) further coupled EPIRICE with geographic information system (GIS) data to conduct multiscale simulations of rice leaf blast and ShB epidemics, enabling effective prediction of disease dynamics across different spatial and temporal scales. These studies demonstrate the effectiveness of mechanistic models in interpretability and their ability in representing disease dynamics. However, the performance of mechanistic models often depends heavily on predefined assumptions and parameter settings (D’Agostino McGowan et al., 2021). These assumptions may limit the model’s adaptability and predictive capability under complex and heterogeneous environmental conditions (Cunniffe et al., 2015; Heesterbeek et al., 2015).

More recently, machine learning (ML), particularly deep learning (DL), has gradually become a dominant paradigm for crop disease forecasting (Dokic et al., 2020; Delfani et al., 2024). Traditional ML algorithms, such as random forests (RF), support vector machines (SVM), and artificial neural networks (ANN), have demonstrated strong capabilities in modelling nonlinear relationships between environmental variables and disease occurrence (Mehra et al., 2016; Fenu and Malloci, 2019; Wadhwa and Malik, 2024). For instance, Wadhwa and Malik (2024) applied a CatBoost model for rice pest forecasting in India, achieving high predictive accuracy across multiple pest categories. Deep learning approaches have further improved the capability to model complex temporal dependencies, particularly in sequential forecasting tasks. Chen et al. (2025) developed long short-term memory (LSTM) and convolutional recurrent hybrid models for tobacco disease forecasting, achieving short-term and long-term prediction *R*^2^ ∼ 0.8. In potato late blight forecasting, Abderrahmane et al. (2022) proposed an LSTM-autoencoder (LSTM-AE) framework capable of predicting outbreaks up to 30 days in advance. Lee and Yun (2023) constructed a deep learning-based forecasting framework using large-scale multi-crop environmental datasets, achieving an average area under the receiver operating characteristic curve (AUROC) of 0.917 across multiple crop pests and diseases. Despite their high predictive performance, these data-driven models are often regarded as black-box systems, lacking explicit biological interpretability. Their generalization capability and predictive reliability may therefore deteriorate substantially under unseen environmental conditions or scenarios not represented in the training data (Fenu and Malloci, 2021; Delfani et al., 2024).

Advances in sensing technologies have substantially expanded the availability of habitat-related information for disease forecasting. Early disease forecasting studies mainly relied on meteorological observations and field survey data collected at discrete spatial locations and time points, limiting continuous characterization of disease-conducive environments. In recent years, the rapid development of remote sensing and Internet of Things (IoT) technologies has facilitated the integration of multi-source observational data into disease forecasting studies, enabling high-resolution spatiotemporal monitoring of crop canopy structure, soil conditions, and microclimatic environments (Jensen et al., 2025). In particular, the capability of remote sensing technologies for large-scale and continuous observation has provided new opportunities for characterizing disease habitat conditions and their spatiotemporal heterogeneity, thereby supplying more comprehensive and continuous inputs for disease forecasting models (Araújo et al., 2021).

The increasing availability of multi-source data has created new opportunities for improving crop disease forecasting. However, effectively integrating heterogeneous environmental information while maintaining biological interpretability remains a major challenge. Given the complementary strengths and limitations of mechanistic and data-driven approaches, hybrid forecasting frameworks integrating multi-source data with epidemiological mechanisms have emerged as a promising solution for addressing this challenge (Karpatne et al., 2017; Reichstein et al., 2019). In this context, incorporating epidemiological knowledge into data-driven models provides an effective strategy for bridging these two paradigms. Specifically, differential equations describing physical or biological processes can be embedded into deep learning frameworks as mechanistic constraints, allowing the models to fit observational data while simultaneously conforming to underlying epidemiological dynamics. Such integration has the potential to improve model robustness, interpretability, and generalization capability. In recent years, physics-informed learning frameworks, which integrate physical or biological mechanisms into deep learning models through governing equations or mechanistic constraints, have attracted increasing attention across multiple scientific domains (Raissi et al., 2019; Cuomo et al., 2022). However, their applications in complex agroecosystems, especially for crop pest and disease forecasting, remain relatively limited.

Given these knowledge gaps, we focus on large-scale spatiotemporal forecasting of ShB and investigate the integration of mechanistic information with data-driven learning. The main objectives of this study are: (1) To construct a multi-source disease habitat characterization framework by integrating remote sensing, meteorological, and soil property data; (2) To develop a physics-informed hybrid gated recurrent unit (PI-HGRU) model by incorporating epidemiological mechanisms; (3) To establish a sliding-window-based temporal forecasting framework for representing cumulative environmental effects and delayed disease responses; and (4) To evaluate the predictive performance of the proposed framework using multi-regional and multi-year disease observation datasets.

## 2. Materials and Methods

The overall workflow of this study is illustrated in Fig. 1, which mainly consists of four components: (a) Data acquisition and preprocessing, (b) Feature engineering and selection, (c) Model construction and training, and (d) Model performance and analysis.

**Fig. 1.**
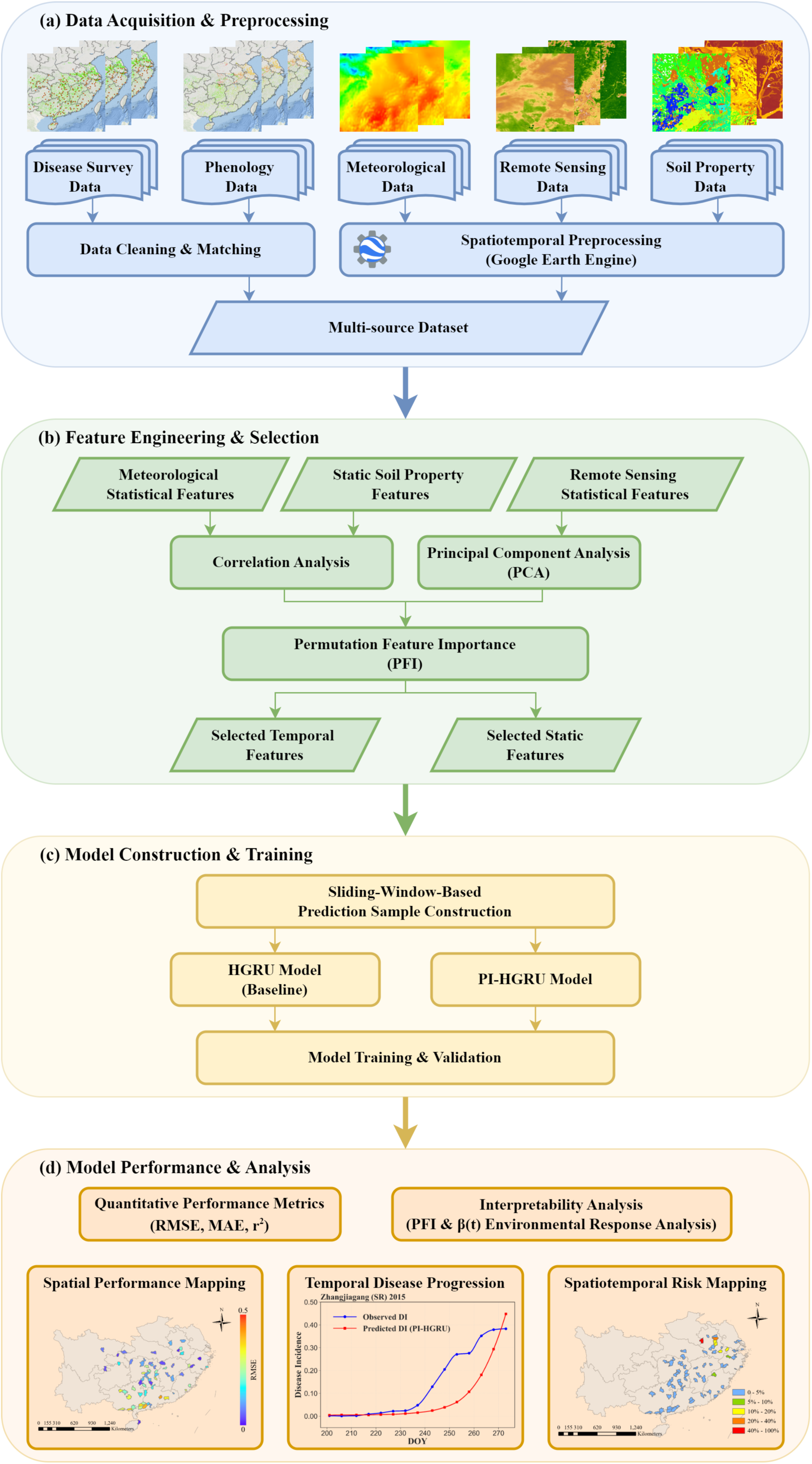
Workflow used in this work to develop a framework for forecasting rice sheath blight (ShB) at large spatiotemporal scales. (a) Disease survey and phenological data are cleaned and matched to construct growing-season disease sequences. Meteorological, remote sensing, and soil property data are then processed and integrated to form the multi-source dataset. (b) Meteorological and remote sensing time series are next summarized as statistical features, while soil property variables are treated as static features in our framework. Correlation analysis and principal component analysis are then used to reduce redundancy and feature dimensionality. Permutation feature importance is then used to identify informative temporal and static variables. (c) The selected variables from the previous step are then organized into prediction samples using the sliding-window-based sample construction framework developed in this study. The resulting samples are used to train and validate the hybrid gated recurrent unit (HGRU) baseline and the proposed physics-informed hybrid gated recurrent unit (PI-HGRU) model. (d) Finally, model performance is evaluated using root mean square error (RMSE), mean absolute error (MAE), and squared Pearson correlation coefficient (*r*²), together with permutation feature importance (PFI) and the learned transmission rate *β*(*t*) for model interpretation. Our trained framework is used for spatial performance mapping, temporal disease progression, and spatiotemporal risk mapping.

### 2.1. Study Area and Disease Observation Data

#### 2.1.1. Study Area

The study area encompasses major rice-growing regions across 16 provinces in southern China (Fig. 2). These regions are primarily located within subtropical and tropical monsoon climate zones, where warm and humid conditions favor the occurrence and development of rice sheath blight (ShB). The broad geographic extent also encompasses substantial variation in climatic and environmental conditions across rice-growing regions.

**Fig. 2.**
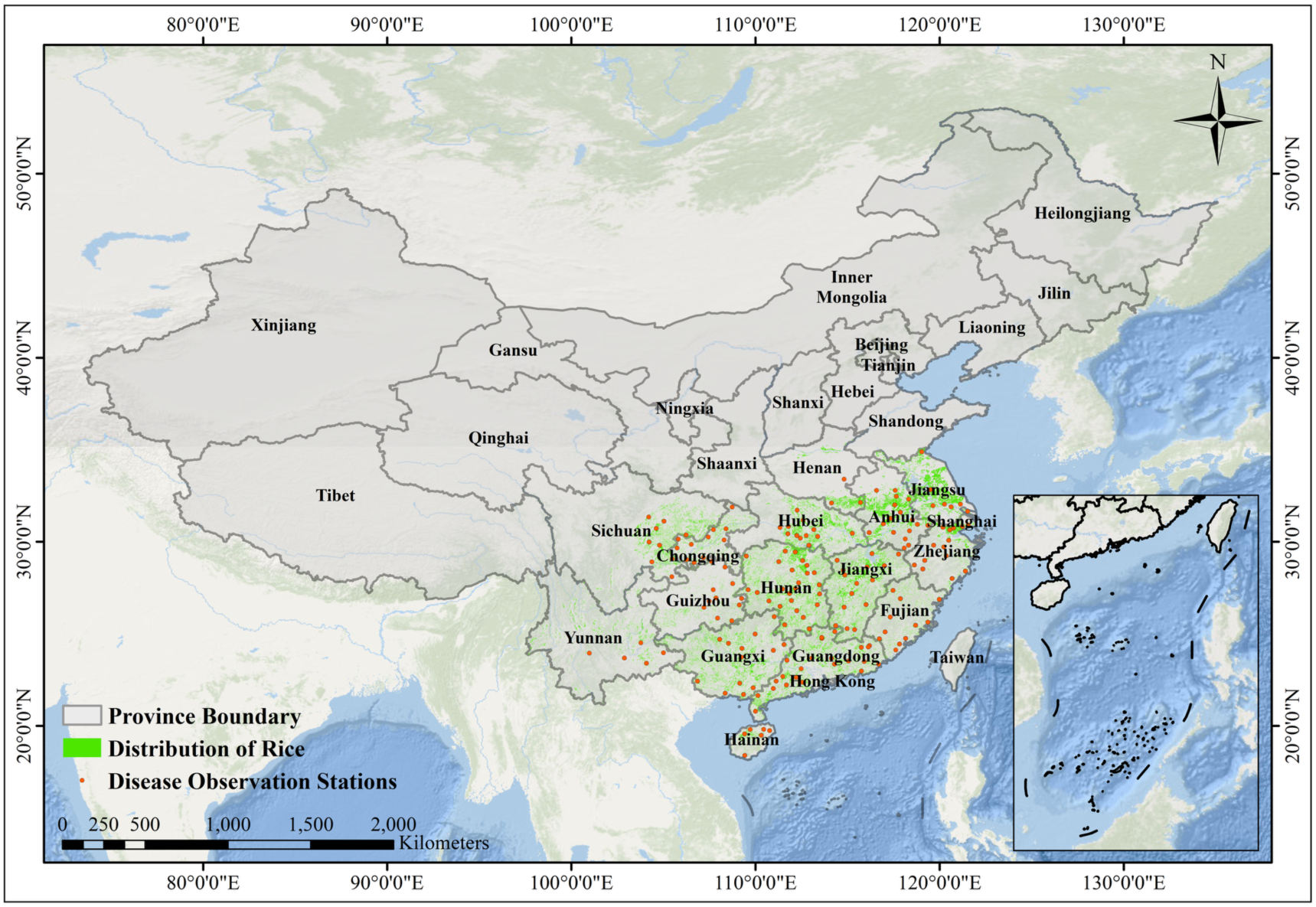
Geographic distribution of the study area, rice-growing regions, and rice sheath blight (ShB) observation stations. The study area covers major rice-growing regions across 16 provinces in southern China. Green areas indicate the spatial distribution of rice cultivation, while orange points represent field observation stations used for ShB monitoring. Provincial boundaries provide geographic context for the spatial distribution of the study area and observation network.

#### 2.1.2. Disease Observation Data and Preprocessing

ShB field surveys were conducted by plant protection specialists from the National Agro-Tech Extension and Service Center (NATESC) from 2000 to 2016. The surveys followed the national standard Rules for Investigation and Forecast of the Rice Sheath Blight (*Rhizoctonia solani* Kühn) (GB/T 15791–1995). The records included observation station locations, survey dates, rice cropping types, phenological stages, and disease incidence rates.

To standardize the disease observations, two preprocessing procedures were applied. (1) Growing-season sequence screening: Each unique combination of observation station, year, and rice cropping type was defined as an individual growing-season sequence. Sequences with fewer than four valid disease incidence observations were excluded because they provided insufficient information to characterize disease progression. (2) Phenological information matching: The retained disease observations were spatially and temporally matched with the *ChinaCropPhen1km* dataset, a 1-km^2^ phenological dataset of the three major grain crops in China from 2000 to 2019 (Luo et al., 2020a). Phenological information was first extracted according to the geographic coordinates of each observation station. A 3 × 3 neighboring window centered on each station was used to identify the nearest valid pixel corresponding to the same rice cropping type, thereby reducing potential mismatches caused by grid-scale heterogeneity. Disease observations were then matched with phenological records for the corresponding year and rice cropping type. The day of year (DOY) values of transplanting (TR), heading (HE), and maturity (MA) were assigned to each growing-season sequence as standardized phenological references. After preprocessing, 1,620 growing-season sequences were retained for subsequent analysis.

To provide an independent evaluation of model performance on later years, the dataset was chronologically divided into training, validation, and test sets. Data from 2000–2011 were used for model training and feature selection, while data from 2013–2014 were used for validation and hyperparameter selection. Data from 2015–2016 were reserved as an independent test set for final performance evaluation. This chronological partition was maintained throughout model development to prevent information leakage from the test set.

### 2.2. Environmental Variables and Data Preprocessing

To construct a comprehensive set of input features for disease forecasting, this study integrated multi-source remote sensing, meteorological, and soil property datasets. The environmental variables and their corresponding data sources are summarized in Table 2. All geospatial preprocessing procedures, including spatial resolution harmonization and regional-scale statistical aggregation, were conducted on the Google Earth Engine (GEE) cloud platform (Gorelick et al., 2017). This processing ensured consistent spatial integration across the geospatial datasets.

**Table 2.**
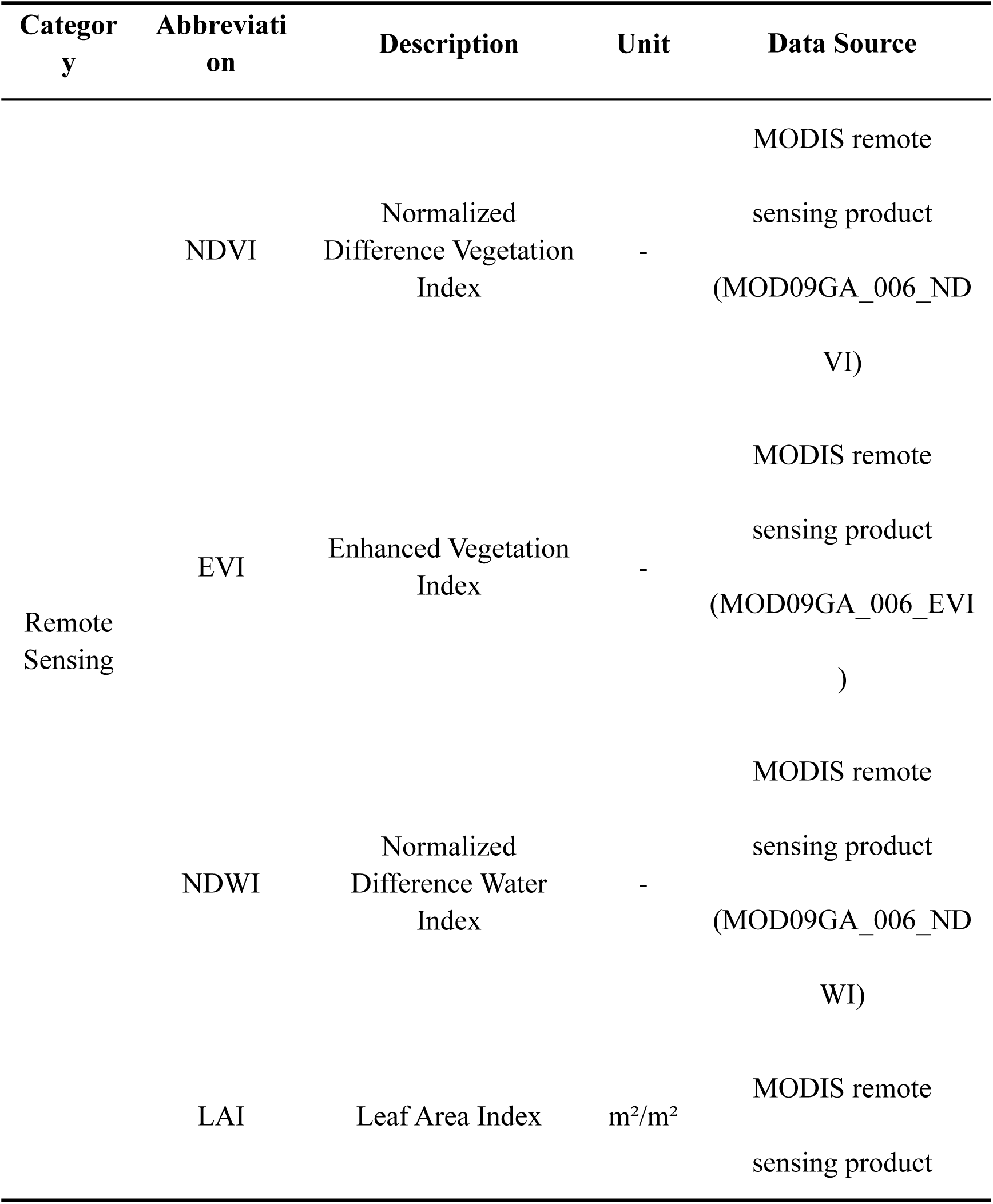
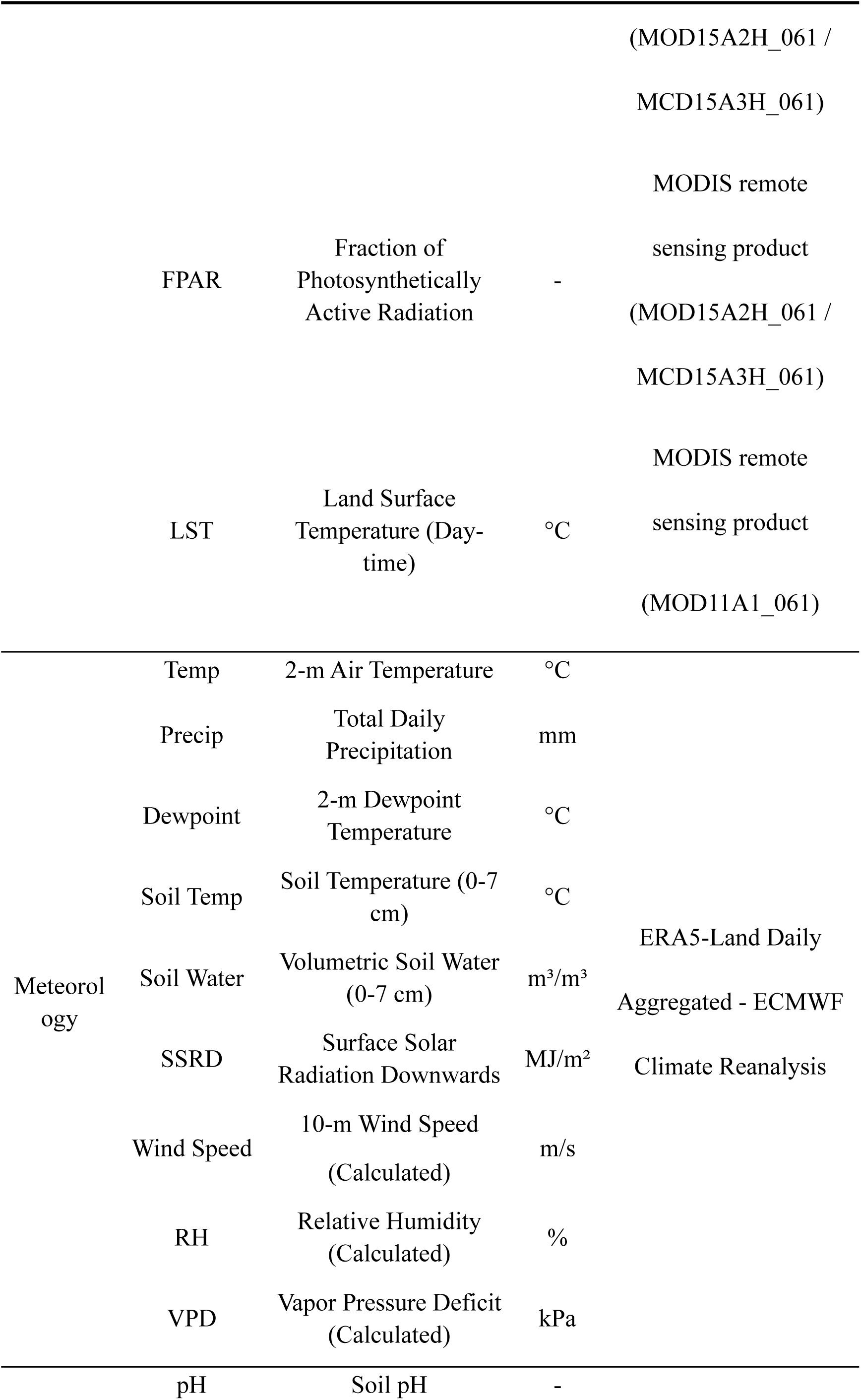
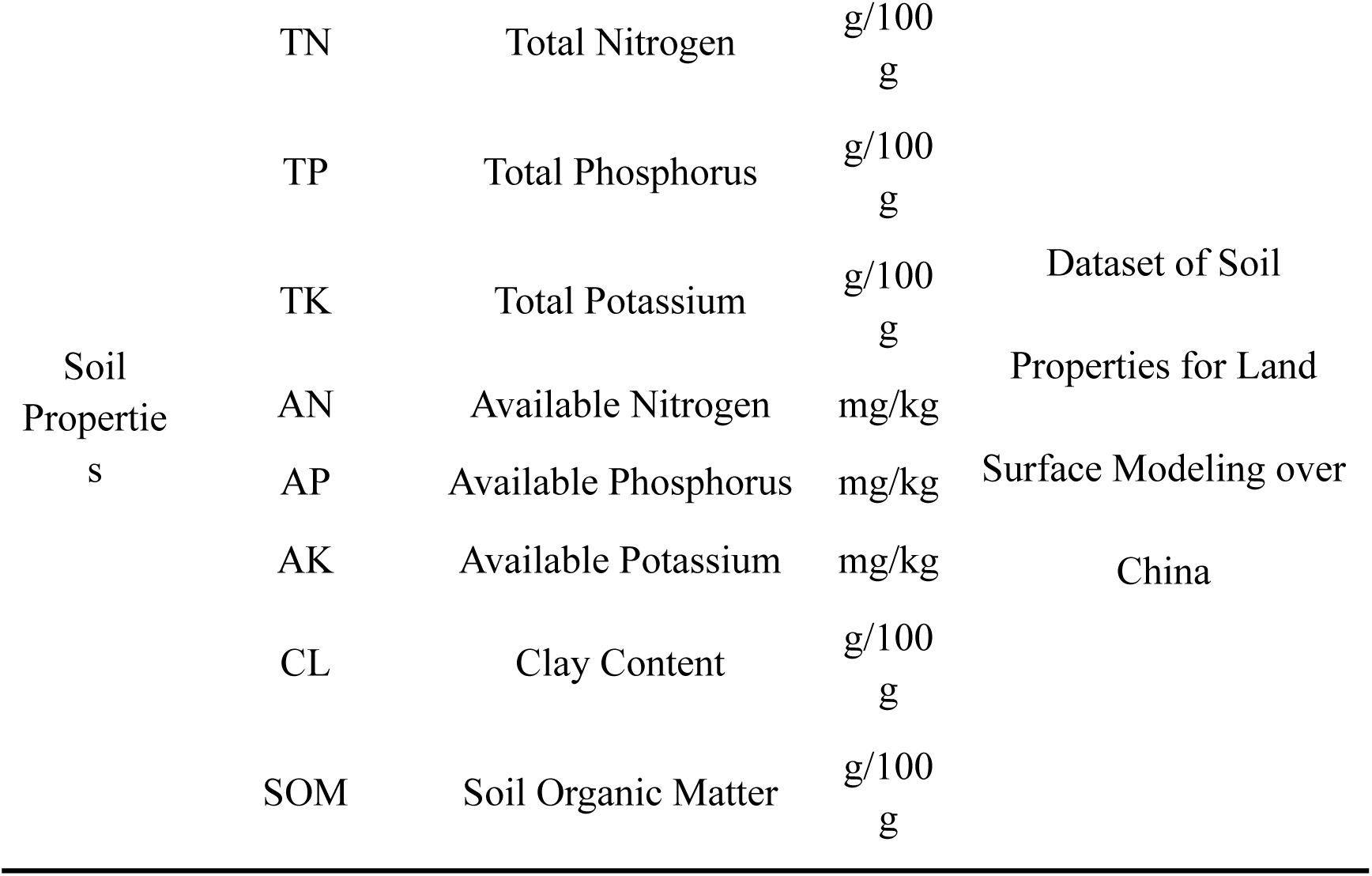
Environmental variables used in this study to forecast rice sheath blight in China. Abbreviations correspond to the variable names used throughout this study. Units refer to values after unit conversion where applicable.

#### 2.2.1. Remote Sensing Data

Remote sensing data were used to characterize crop growth conditions and environmental dynamics associated with disease development. The datasets were obtained from Moderate Resolution Imaging Spectroradiometer (MODIS) products distributed by the NASA Land Processes Distributed Active Archive Center (LP DAAC). Three categories of MODIS products were used. First, leaf area index (LAI) and fraction of photosynthetically active radiation (FPAR) were obtained from MOD15A2H Version 6.1 (500 m, 2000–2002) and MCD15A3H Version 6.1 (500 m, 2003–2016). These products provide information on vegetation structural conditions. Second, normalized difference vegetation index (NDVI), enhanced vegetation index (EVI), and normalized difference water index (NDWI) were calculated from the MODIS/Terra surface reflectance product MOD09GA Collection 6. Third, daily land surface temperature (LST) was obtained from MOD11A1 Version 6.1 (1 km) to characterize surface thermal conditions.

For data preprocessing, annual rice cultivation maps were obtained from the ChinaCropArea1km dataset, which provides 1-km² distributions of three major grain crops in China from 2000 to 2019 (Luo et al., 2020b). These maps were used to identify rice-growing areas within each county for the corresponding year. Quality control was then conducted using the quality assessment (QA) layers provided with each MODIS product. Only observations classified as high or moderate quality were retained. Observations affected by cloud contamination, cloud shadows, or high aerosol concentrations were excluded. All valid observations within rice-growing areas of each county were spatially averaged to generate county-scale remote sensing variables.

To harmonize temporal resolutions among MODIS products and improve temporal continuity, time-series reconstruction was performed for all remote sensing variables. Missing observations were interpolated using the Akima spline method to generate continuous daily time series (Akima, 1970). Akima interpolation limits excessive oscillations while maintaining curve smoothness, making it suitable for reconstructing irregularly spaced observations. Interpolation was restricted to the temporal range of available observations, without extrapolation.

#### 2.2.2. Meteorological Data

Meteorological conditions, particularly temperature- and moisture-related variables, are among the most direct environmental drivers governing the occurrence and progression of fungal diseases such as ShB (Huber and Gillespie, 1992; Bhukal et al., 2015). Accordingly, to characterize the meteorological drivers of ShB development, daily meteorological data were obtained from the land component of the fifth-generation European Centre for Medium-Range Weather Forecasts (ECMWF) reanalysis dataset (ERA5-Land) (Muñoz-Sabater et al., 2021). The data were accessed through the ERA5-Land Daily Aggregated product available on the GEE platform, which provides spatially and temporally continuous meteorological information. The ERA5-Land variables used in this study include daily mean 2-m air temperature, total daily precipitation, daily mean 2-m dewpoint temperature, and daily mean soil temperature and volumetric soil water in the 0–7 cm surface layer. Daily accumulated surface solar radiation downwards (SSRD) and daily mean eastward (*u*) and northward (*v*) wind components at 10 m height were also included. Relative humidity (RH), vapor pressure deficit (VPD), and 10-m wind speed were derived from these original meteorological variables. All ERA5-Land grid cells within each county were spatially averaged to represent county-scale meteorological conditions.

RH and VPD were calculated from daily mean air temperature (*T*) and dewpoint temperature (*T_d_*) using the August–Roche–Magnus approximation (Bolton, 1980; Lawrence, 2005). Saturation vapor pressure (*e_s_*) and actual vapor pressure (*e*) were first estimated using Eqs. (1) and (2):

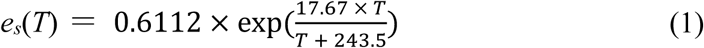

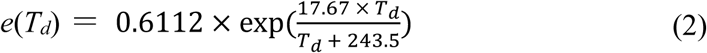

where *e_s_*(*T*) denotes saturation vapor pressure calculated from air temperature *T*, and *e*(*T_d_*) denotes actual vapor pressure calculated from dewpoint temperature *T_d_*. Both vapor pressures are expressed in kPa, while *T* and *T_d_* are expressed in °C. RH (%) and VPD (kPa) were then calculated using Eqs. (3) and (4), respectively:

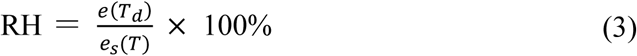

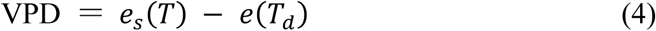

In addition, the daily mean 10-m wind speed (m/s) was calculated from the eastward (u) and northward (v) wind components using Eq. (5):

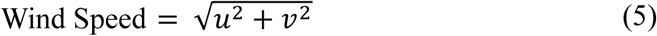

#### 2.2.3. Soil Property Data

Soil physicochemical properties may influence disease occurrence and development through their relationships with crop growth, root-zone conditions, and field moisture dynamics (Janvier et al., 2007; Berendsen et al., 2012). Soil property data were obtained from the Dataset of Soil Properties for Land Surface Modeling over China, which provides multilayer soil physicochemical attributes at a spatial resolution of 1 km across China (Shangguan et al., 2013). Because soil properties are relatively stable over time, these variables were incorporated as static environmental features. Soil variables within the rice-growing areas of each county were spatially averaged to represent county-scale conditions. The candidate soil variables included soil organic matter (SOM), soil pH, total nitrogen (TN), total phosphorus (TP), total potassium (TK), available nitrogen (AN), available phosphorus (AP), available potassium (AK), and clay content (CL). These variables characterize soil nutrient status, physicochemical properties, and texture conditions relevant to crop growth and disease development.

### 2.3. Feature Selection and Modelling Data Preparation

#### 2.3.1. Feature Selection Strategy

To reduce input-feature dimensionality, alleviate multicollinearity, and identify informative environmental predictors for disease forecasting, a two-stage selection strategy was used. The first stage focused on dimensionality reduction and redundancy elimination within each data category (Chandrashekar and Sahin, 2014). Daily meteorological variables within each growing season were summarized using the mean, standard deviation, median, minimum, maximum, and coefficient of variation, while soil property variables were treated as static features. Pearson correlation matrices were calculated separately for meteorological statistical features and soil property variables (Figs. S1 and S2), and highly correlated features (|*r*|, 0.90) were removed while retaining representative environmental information (Smith et al., 2024). Strong correlations between Temp and Soil Temp, RH and VPD, and TN and SOM led to the removal of Soil Temp, RH, and TN, respectively. The median and coefficient of variation were also excluded because of their redundancy with other statistical features.

Remote sensing variables were summarized using the same growing-season statistics and further reduced using principal component analysis (PCA) because of substantial information overlap among the resulting features (Harris and Van Niekerk, 2018; Kiala et al., 2020). The statistical features were transformed into mutually orthogonal principal components (PCs), and PCs explaining more than 95% of the cumulative variance were retained (Jolliffe and Cadima, 2016). This procedure reduced the 36 remote sensing statistical features to 14 PCs.

The second stage involved global feature importance evaluation across all feature categories. The retained meteorological features, soil property variables, and remote sensing PCs were used to construct a random forest regression model, with the area under the disease progress curve (AUDPC) as the target variable. Permutation feature importance (PFI) was calculated on the validation dataset (Breiman, 2001), and features within the top 50% of the PFI ranking were retained for subsequent model construction. Meteorological and soil features were traced directly to their original variables. For remote sensing PCs, the contribution of each original remote sensing variable was quantified by aggregating its squared loadings across the highly ranked PCs.

#### 2.3.2. Construction of Prediction Samples Based on a Sliding Time Window

To fully exploit the temporal information contained in environmental variables, we designed a sliding-window-based sample construction framework. This approach was needed because disease development is not an instantaneous response to environmental changes. Rather, disease development reflects the cumulative effects of preceding environmental conditions and the delayed responses associated with pathogen infection, symptom development, and subsequent disease progression (Biswas et al., 2011; Pal et al., 2017). Therefore, two temporal parameters, namely the input time window (*T_input_*) and lag time (*T_la_*_9_), were introduced to characterize these processes during sample construction.

The two temporal parameters were jointly used to construct standardized prediction samples for each disease observation. As shown in Fig.3, *T_input_* defines the length of historical environmental information used for prediction, while *T_la_*_9_ defines the interval between the end of the environmental input window and the target disease observation. By jointly introducing these two parameters, the constructed samples represent both cumulative environmental conditions and lagged disease responses. Specifically, for each growing-season sequence, every disease observation *Y*(*t*) was treated as an independent prediction target (*Y_tar_*_9*et*_), where *Y*(*t*) denotes the observed disease incidence at day of year *t*. Daily remote sensing and meteorological variables within the period 8*t* − *T_la_*_9_ − *T_input_*, *t* − *T_la_*_9_ − 1: were extracted as temporal input features (*X_temporal_*). This time window represents the historical environmental conditions preceding the target disease observation while accounting for the lagged response of disease development to environmental drivers. Together with the static soil property variables (*X_static_*), a complete prediction sample *S* can be expressed as Eq. (6):

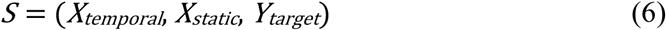

**Fig. 3.**
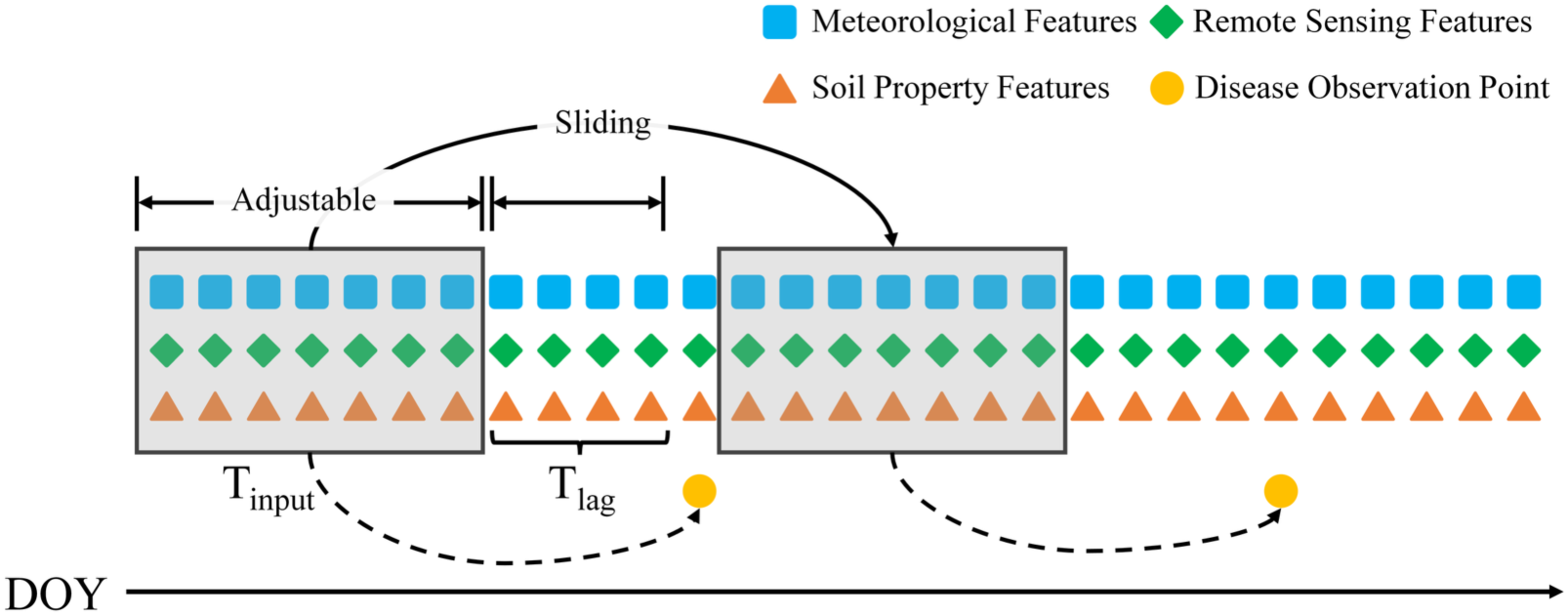
Schematic illustration of prediction sample generation using the sliding-window-based framework developed in this study. *T_input_* defines the adjustable length of the historical environmental input window, while *T_la_*_9_ defines the adjustable interval between the end of this window and the target disease observation point. Meteorological and remote sensing features within the historical window are used as temporal inputs, while soil property features are treated as static inputs. These inputs are paired with the corresponding rice sheath blight observation to construct a prediction sample. The window then slides along the day-of-year sequence to generate samples for subsequent model training and evaluation.

By applying the aforementioned sliding-window sampling procedure to all observation time points within each growing-season sequence, a structurally consistent prediction sample dataset was generated for subsequent model training and evaluation. The resulting samples preserve the temporal relationships between environmental conditions and disease development, while providing a unified input format for spatiotemporal disease forecasting.

### 2.4. Prediction Model Construction and Experimental Settings

#### 2.4.1. Physics-Informed Hybrid Gated Recurrent Unit (PI-HGRU) Model

To address the limited interpretability and generalization of conventional data-driven models, this study developed a physics-informed hybrid gated recurrent unit (PI-HGRU) model by integrating deep learning–based representation learning with epidemiological constraints. The proposed framework combines multi-source environmental information with disease epidemic dynamics within a unified forecasting model. As shown in Fig. 4, PI-HGRU comprises a shared feature encoder, a hybrid prediction module, and a dynamic fusion module. The Shared Feature Encoder processes heterogeneous inputs through two branches that separately encode temporal and static features. The Temporal Encoder consists of a gated recurrent unit (GRU) network combined with a self-attention mechanism to encode the temporal feature sequence *X_temporal_* of length *T_input_*, thereby capturing temporal dependencies within the historical environmental information. The Static Encoder uses a multilayer perceptron (MLP) to learn representations from the static soil features *X_static_*. The outputs of the two encoders are concatenated to form a fused feature vector (Fused Vector), which provides the shared representation for the subsequent prediction modules. Dropout regularization is applied to the fused representation before it is passed to the data-driven and physics-informed heads.

**Fig. 4.**
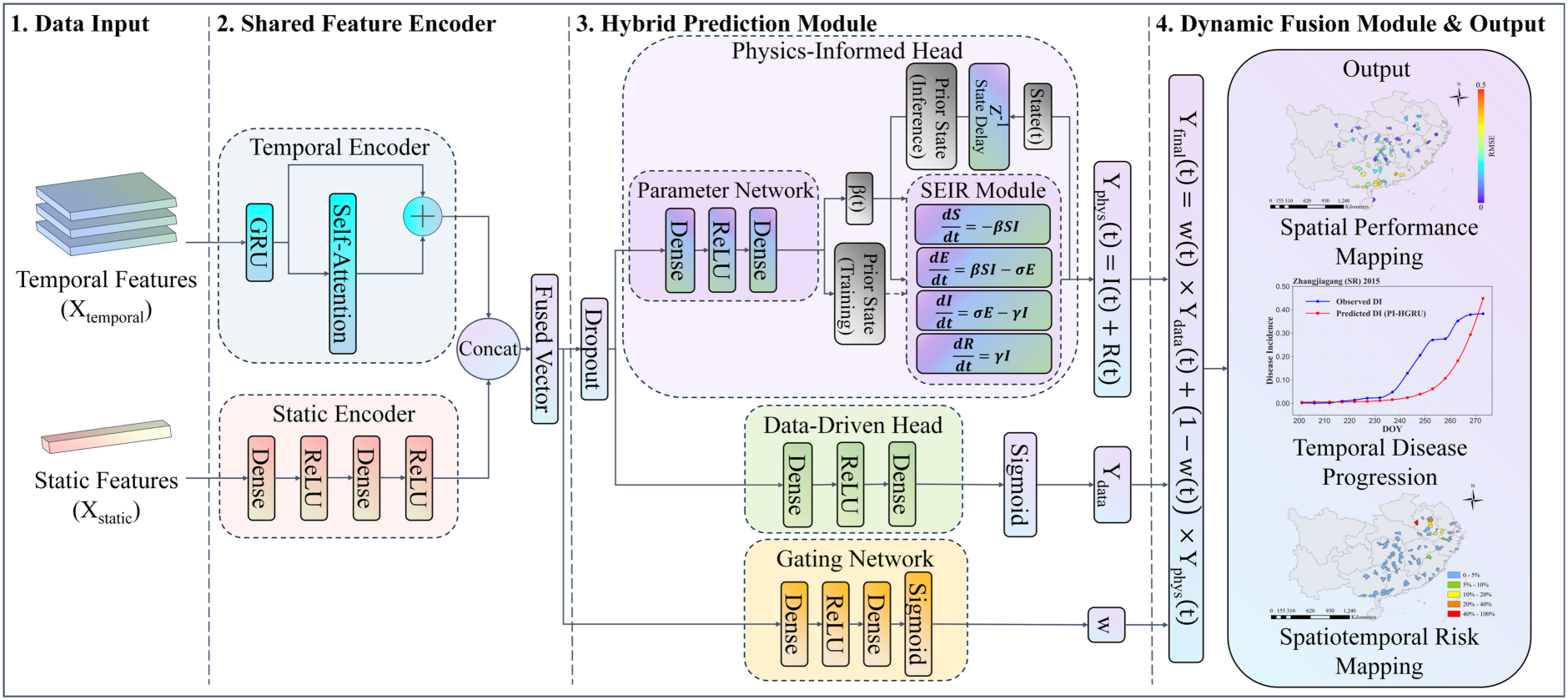
Architecture of the proposed physics-informed hybrid gated recurrent unit (PI-HGRU) model. Temporal environmental features and static soil property features are provided as separate inputs. The shared feature encoder processes temporal inputs using a gated recurrent unit (GRU) with self-attention, while static inputs are encoded using a multilayer perceptron.

The Hybrid Prediction Module constituted the core component of the PI-HGRU model and was designed to simultaneously capture epidemiological dynamics and complex nonlinear relationships based on the shared feature representation. This module consisted of three components: a physics-informed head, a data-driven head, and a gating network.

1. Physics-Informed Head: This component incorporated the susceptible–exposed– infectious–removed (SEIR) epidemiological framework (Hethcote, 2000) into the forecasting process through mechanistic constraints, while its key epidemiological parameters were dynamically estimated by neural networks. Specifically, the fused feature representation was fed into a parameter network to estimate the time-varying infection rate *β*(*t*). During training, the parameter network also provided the prior epidemiological state required for the SEIR update, whereas during inference the updated SEIR state was propagated between successive prediction steps. This design allowed the infection rate to vary dynamically with environmental conditions while maintaining the epidemiological state information required for mechanistic prediction. The latent transition rate *σ* and removal rate *γ* were fixed at *σ* = 1/3 and *γ* = 1/70, respectively, while the parameter network for *β*(*t*) was initialized with a reference value of 0.46. These settings were informed by previous epidemiological studies (Rodrigues et al., 2003; Kim et al., 2015). Given the estimated *β*(*t*) and the corresponding prior epidemiological state, the SEIR states were updated according to the disease transmission dynamics described in Eq. (7):

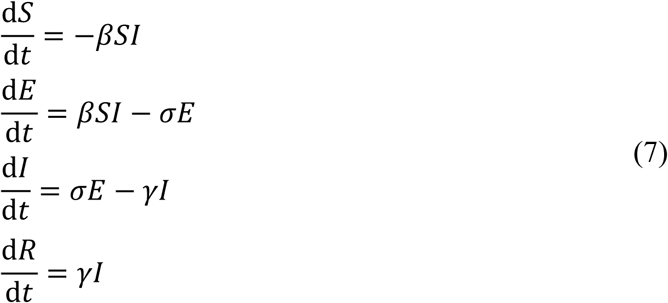

where *S*, *E*, *I*, and *R* denote the proportions of susceptible, exposed, infectious, and removed plants, respectively (Hethcote, 2000); *β* represents the infection rate; *σ* denotes the transition rate from the exposed to the infectious state; and *γ* denotes the transition rate from the infectious to the removed state. The equations were discretized using a one-step Euler integration scheme to update the epidemiological states from the prior state to the current state. To maintain consistency with the field-observed disease incidence, the physics-informed prediction *Y_p_*_ℎ*ys*_(*t*) was calculated as the sum of infectious and removed plant proportions.
2. Data-Driven Head: This component directly learns complex nonlinear relationships between multi-source environmental features and disease development from data, thereby providing complementary predictive information beyond the epidemiological mechanism. Specifically, the fused feature vector was fed into an MLP to generate the data-driven prediction *Y_data_*(*t*), with a sigmoid activation function used to constrain the output to the normalized range of disease incidence. By leveraging the expressive power of deep learning, this branch can model complex environmental influences and latent patterns that are difficult to explicitly represent using mechanistic equations alone.
3. Gating Network: To achieve adaptive fusion between the physics-informed prediction and the data-driven prediction, a learnable gating mechanism was introduced. Taking the fused feature vector as input, the gating network generated a dynamic fusion weight *w*(*t*) ∈ [0, 1], which adjusted the relative contributions of the two prediction branches according to the input conditions. This design allowed the model to adaptively balance the data-driven and physics-informed predictions rather than assigning a fixed contribution to either branch.

The Dynamic Fusion Module generated the final prediction by adaptively combining the outputs of the data-driven and physics-informed branches according to the dynamic fusion weight *w*(*t*), as formulated in Eq. (8):

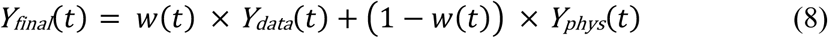

where *Y_final_*(*t*) denotes the final prediction output at time *t*; *Y_data_*(*t*) and *Y_p_*_ℎ*ys*_(*t*) represent the outputs generated by the data-driven head and the physics-informed head, respectively; and *w*(*t*) ∈ [0,1] denotes the dynamic fusion weight produced by the gating network. This mechanism enables the model to adaptively integrate data-driven learning and mechanistic constraints according to the characteristics of the input conditions, thereby improving the forecasting robustness and generalization capability.

To assess the contribution of physics-informed constraints, a hybrid gated recurrent unit (HGRU) model was constructed as the data-driven baseline based on generic hybrid sequential modelling frameworks (Lim et al., 2021; Gao et al., 2023), with its architecture adapted to the prediction task and input data structure used in this study. Structurally, HGRU retained the same temporal and static feature encoders and the same data-driven prediction head as PI-HGRU, while excluding the physics-informed head, gating network, and dynamic fusion module. Both models therefore used the same input features and shared the same data-driven feature representation and prediction architecture, and were trained and evaluated under identical experimental settings. This controlled comparison allowed the contribution of introducing the physics-informed components to model performance to be evaluated while minimizing differences arising from other architectural or experimental factors.

The resulting representations are concatenated into a fused feature vector, followed by dropout before entering the two prediction heads. The hybrid prediction module contains a physics-informed head, a data-driven head, and a gating network. The physics-informed head estimates the time-varying transmission rate *β*(*t*) and updates the susceptible–exposed–infectious– removed epidemiological states to generate the physics-informed prediction *Y_p_*_ℎ*ys*_(*t*). In parallel, the data-driven head generates *Y_data_*(*t*) directly from the fused environmental representation. The gating network produces a dynamic weight *w*(*t*), which adaptively combines *Y_data_*(*t*) and *Y_p_*_ℎ*ys*_(*t*) to obtain the final prediction *Y_final_*(*t*). The final outputs are used to characterize spatial forecasting performance, temporal disease progression, and spatiotemporal rice sheath blight risk.

#### 2.4.2. Model Training and Evaluation Strategy

A time window sensitivity analysis was conducted to determine the input time window and lag time within the sliding time-window forecasting framework. This step was carried out using the HGRU model. Different combinations of *T_input_* and *T_lag_* were evaluated on the validation dataset, and the best-performing configuration was fixed for subsequent model training and comparative experiments.

During model training, all models were optimized using the Adam optimizer. The learning rate was set to 1 × 10^-5^, the batch size to 64, and the maximum number of training epochs to 200. To mitigate overfitting, early stopping was applied, with training terminated if the validation loss did not improve for 15 consecutive epochs. For PI-HGRU, training was guided by a composite loss function consisting of a data loss term and a mechanistic consistency loss term. The overall objective function was formulated as Eq. (9):

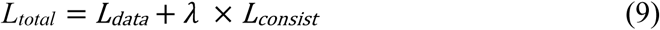

where *λ* denotes the weighting coefficient of the consistency loss and was set to 0.5 based on validation-set performance. The data loss term *L_data_* was implemented using Smooth L1 Loss to measure the discrepancy between the final fused prediction *Y_final_* and observed disease incidence. The consistency loss *L_consist_* was defined as the mean squared error (MSE) between the predictions generated by the data-driven head (*Y_data_*) and the physics-informed head (*Y_p_*_ℎ*ys*_). This term encouraged consistency between the two prediction branches and served as a regularization mechanism during training. For the HGRU baseline model, only the data loss term *L_data_* was used as the training objective. Considering the temporal discontinuity of disease observations, the data loss was computed only at time points with valid observations.

Model performance was evaluated using the independent test dataset. Within each growing-season sequence, step-wise rolling forecasting was performed across successive prediction targets. At each prediction step, the model used the corresponding historical environmental sequence and static soil features, while PI-HGRU propagated the latent SEIR states from the preceding prediction target to preserve epidemiological continuity. Model performance was evaluated using the RMSE, mean absolute error (MAE), and squared Pearson correlation coefficient (*r*^2^), as formulated in Eqs. (10) – (12), respectively.

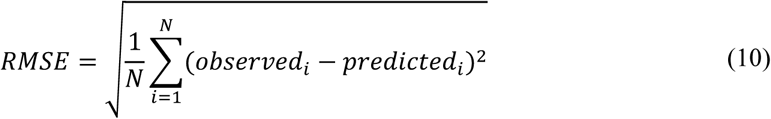

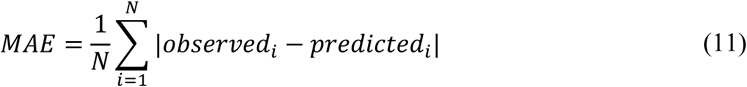

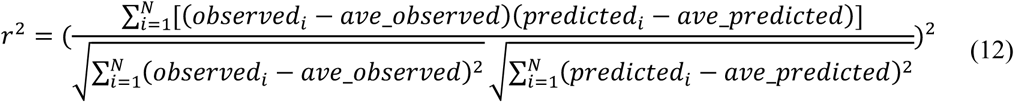

where *N* denotes the number of valid observation points within a growing-season sequence; *observed_i_* and *predicted_i_* represent the temporally matched observed and predicted values at the *i*-th observation point, respectively; and *ave*_*observed* and *ave*_*predicted* denote their corresponding mean values. Evaluation metrics were calculated only at time points with valid disease observations.

To account for variability due to random initialization, each model was independently trained five times using different random seeds. The mean and standard deviation of the run-level evaluation metrics were reported as the overall model performance. All model training and experiments were conducted on a computational platform equipped with an NVIDIA RTX 4090 GPU (24 GB), a 16-core Intel Xeon Platinum 8352V CPU (2.10 GHz), and 120 GB RAM.

#### 2.4.3. Model Interpretability Analysis

Two complementary analyses were conducted to interpret the environmental information used by PI-HGRU. First, PFI was calculated for a trained PI-HGRU model using the independent test dataset. Each input variable was individually permuted while the remaining variables were kept unchanged, and model predictions were recalculated using the same step-wise rolling forecasting procedure described above. Feature importance was quantified as the increase in mean RMSE across growing-season sequences relative to the unpermuted prediction. Larger increases in RMSE therefore indicated greater contributions of the corresponding variables to model prediction.

Second, the environmental relationships of the learned transmission rate *β*(*t*) were examined. The *β*(*t*) values generated during step-wise rolling forecasting were extracted for all prediction targets in the test dataset. For each of the 13 temporal environmental variables, the mean, standard deviation, minimum, and maximum were calculated within the corresponding historical input window, yielding 52 environmental statistics. Pearson correlation coefficients (*r*) were then calculated between these statistics and the corresponding *β*(*t*) values to characterize their linear associations. These analyses provided complementary perspectives on variable contribution to prediction and environmental relationships with the modeled disease transmission dynamics.

## 3. Results

### 3.1. Input Feature Selection

The second-stage cross-category feature selection identified important predictors from all three data categories (Fig. S3). Minimum vapor pressure deficit (VPD(min)) and the standard deviation of surface solar radiation downwards (SSRD(std)) ranked highly among the meteorological features. Total potassium (TK), total phosphorus (TP), and clay content (CL) were among the most important soil property variables. Several remote sensing principal components (PCs), including RS_PC_8, also showed relatively high permutation feature importance (PFI) scores.

Five remote sensing PCs were included in the top 50% of the PFI ranking. Analysis of their squared loadings showed that all six original remote sensing variables contributed to these components, with overall contributions ranging from 9.9% to 28.4%. Accordingly, all six remote sensing variables were retained for subsequent time-series modelling. Together with the meteorological and soil variables selected by PFI, the final input feature set consisted of 13 temporal variables and eight static variables.

### 3.2. Time Window Sensitivity Analysis

To determine the temporal configuration for model input, a sensitivity analysis was conducted using different combinations of input time window (*T_input_* = 15, 30, 45, and 60 days) and lag time (*T_la_*_9_ = 1, 3, 5, and 7 days). The predictive performance of each configuration was evaluated on the validation dataset using the hybrid gated recurrent unit (HGRU) model (Table 3). Model performance varied across temporal configurations, with no monotonic trend as *T_input_* or *T_la_*_9_ increased. Configurations with *T_input_* = 15 days generally showed poorer performance. Performance improved when *T_input_* increased to 30 or 45 days, whereas extending the window to 60 days provided no consistent improvement. Performance also varied across lag times within each input window. Among the tested configurations, *T_input_* = 45 days and *T_la_*_9_ = 7 days provided the best overall balance across the three metrics. This configuration achieved the lowest root mean square error (RMSE) (0.1370), a near-minimum mean absolute error (MAE) (0.1175), and a squared Pearson correlation coefficient (*r*^2^) of 0.5299. Therefore, a 45-day input window with a 7-day lag was selected for all subsequent experiments.

**Table 3.**
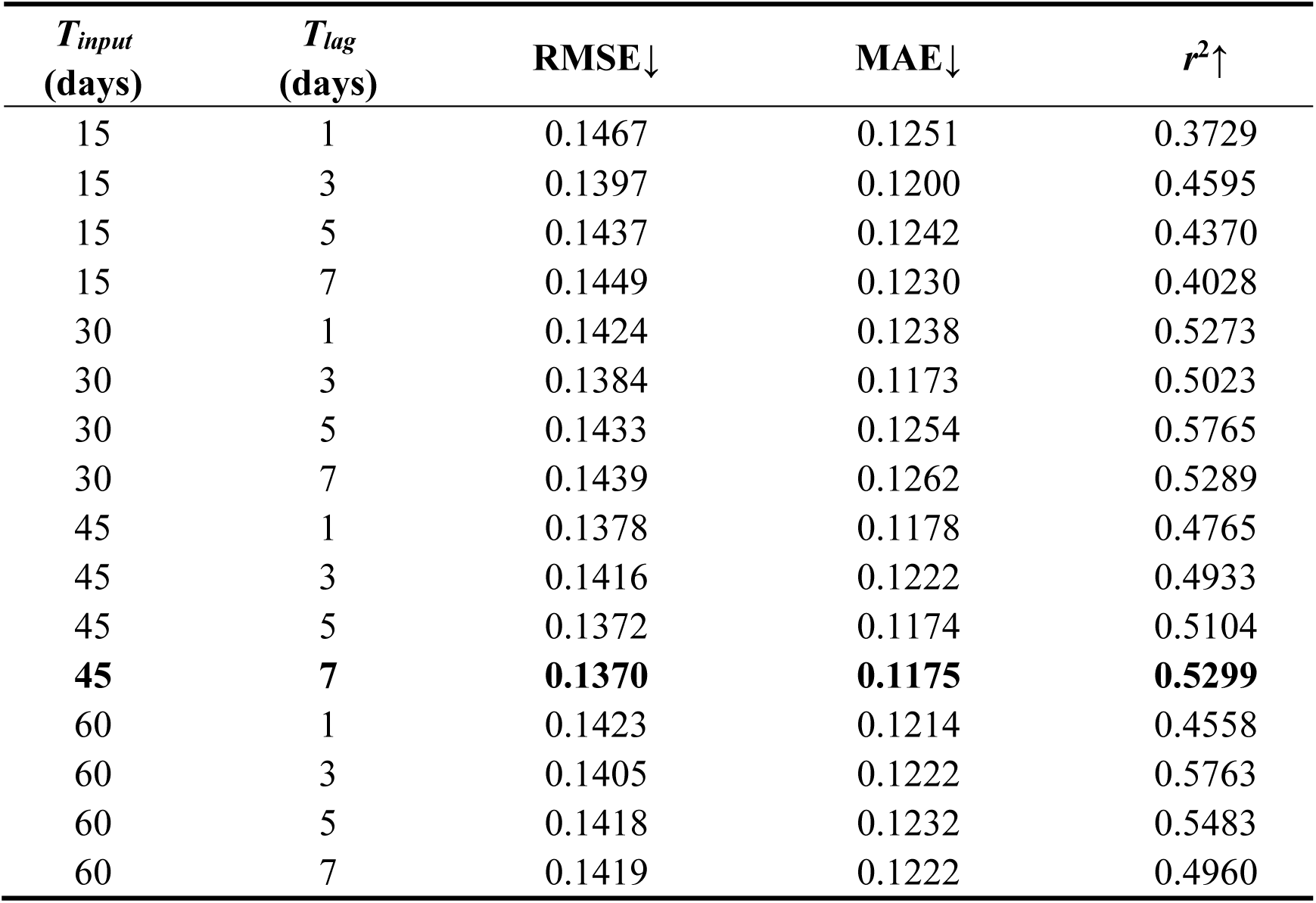
Predictive performance of the hybrid gated recurrent unit (HGRU) model under different input time windows and lag times on the validation dataset. *T_input_* denotes the length of the historical environmental input window, and *T_lag_* denotes the interval between the end of the input window and the target disease observation. Model performance is evaluated using root mean square error (RMSE), mean absolute error (MAE), and squared Pearson correlation coefficient (*r*^2^). ↓ and ↑ indicate that lower and higher values represent better performance, respectively. The selected combination (*T_input_* = 45 days and *T_lag_* = 7 days) is highlighted in bold.

| $T_{input}$<br>(days) | $T_{lag}$<br>(days) | RMSE↓ | MAE↓ | $r^2\uparrow$ |
| --- | --- | --- | --- | --- |
| 15 | 1 | 0.1467 | 0.1251 | 0.3729 |
| 15 | 3 | 0.1397 | 0.1200 | 0.4595 |
| 15 | 5 | 0.1437 | 0.1242 | 0.4370 |
| 15 | 7 | 0.1449 | 0.1230 | 0.4028 |
| 30 | 1 | 0.1424 | 0.1238 | 0.5273 |
| 30 | 3 | 0.1384 | 0.1173 | 0.5023 |
| 30 | 5 | 0.1433 | 0.1254 | 0.5765 |
| 30 | 7 | 0.1439 | 0.1262 | 0.5289 |
| 45 | 1 | 0.1378 | 0.1178 | 0.4765 |
| 45 | 3 | 0.1416 | 0.1222 | 0.4933 |
| 45 | 5 | 0.1372 | 0.1174 | 0.5104 |
| **45** | **7** | **0.1370** | **0.1175** | **0.5299** |
| 60 | 1 | 0.1423 | 0.1214 | 0.4558 |
| 60 | 3 | 0.1405 | 0.1222 | 0.5763 |
| 60 | 5 | 0.1418 | 0.1232 | 0.5483 |
| 60 | 7 | 0.1419 | 0.1222 | 0.4960 |

### 3.3. Evaluation of Model Prediction Performance

The predictive performance of the proposed physics-informed hybrid gated recurrent unit (PI-HGRU) model on the independent test set is summarized in Table 4. Across five independent runs, PI-HGRU achieved an average RMSE of 0.1158, MAE of 0.0958, and *r*² of 0.5751. Compared with the HGRU baseline, PI-HGRU consistently achieved better mean performance across all three metrics. The average RMSE decreased from 0.1290 to 0.1158, corresponding to a reduction of 10.2%. The average MAE decreased from 0.1168 to 0.0958, representing an improvement of 18.0%. Meanwhile, the average *r*² increased from 0.4685 to 0.5751, corresponding to an improvement of 22.8%.

**Table 4.** Predictive performance of the physics-informed hybrid gated recurrent unit (PI-HGRU) model and the hybrid gated recurrent unit (HGRU) baseline across five independent runs on the 2015–2016 test set. Each model is independently trained five times using different random seeds. Mean ± S.D. represents the mean and standard deviation of each evaluation metric across the five runs. RMSE denotes root mean square error, MAE denotes mean absolute error, and *r*² denotes the squared Pearson correlation coefficient. ↓ and ↑ indicate that lower and higher values represent better performance, respectively.

| Model | Seed | RMSE↓ | MAE↓ | $r^2$↑ |
| --- | --- | --- | --- | --- |
| HGRU | Seed1 | 0.1288 | 0.1169 | 0.4643 |
|  | Seed2 | 0.1326 | 0.1203 | 0.4435 |
|  | Seed3 | 0.1241 | 0.1117 | 0.4805 |
|  | Seed4 | 0.1302 | 0.1182 | 0.4856 |
|  | Seed5 | 0.1291 | 0.1169 | 0.4684 |
|  | Mean ± S.D. | **0.1290 ± 0.0031** | **0.1168 ± 0.0032** | **0.4685 ± 0.0164** |
| PI-HGRU | Seed1 | 0.1220 | 0.0988 | 0.5688 |
|  | Seed2 | 0.1039 | 0.0889 | 0.5172 |
|  | Seed3 | 0.1287 | 0.1048 | 0.6708 |
|  | Seed4 | 0.1013 | 0.0867 | 0.5317 |
|  | Seed5 | 0.1230 | 0.0997 | 0.5872 |
|  | Mean ± S.D. | **0.1158 ± 0.0123** | **0.0958 ± 0.0077** | **0.5751 ± 0.0604** |

The performance advantage of PI-HGRU over HGRU was also evident across individual growing-season sequences. The distributions of sequence-level prediction performance across the independent test set are shown in Fig. S4. Each distribution pools the evaluation metrics of individual growing-season sequences across the five independent runs. Compared with HGRU, PI-HGRU showed lower median RMSE and MAE values and a higher median *r*². The *r*² distribution also shifted toward higher values overall. However, the distributions remained broad for both models, indicating substantial variation in predictive performance among growing-season sequences.

### 3.4. Large-Scale Spatiotemporal Disease Forecasting Results

To examine temporal forecasting performance, predicted disease progression curves from PI-HGRU and HGRU were compared with observed values for representative growing-season sequences from the test set (Fig. 5). HGRU showed considerable variation in prediction patterns among the selected sequences. In some cases, its predictions remained relatively stable despite substantial changes in observed disease incidence, whereas in Shaowu it captured some of the observed temporal fluctuations but generally overestimated disease incidence. PI-HGRU showed a more consistent temporal pattern across the selected sequences, with predictions generally increasing as the growing season progressed. This pattern more closely reflected the overall progression of the observed disease incidence. Although discrepancies remained at individual observation dates, PI-HGRU better represented the temporal progression of rice sheath blight (ShB) across the selected sequences.

**Fig. 5.**
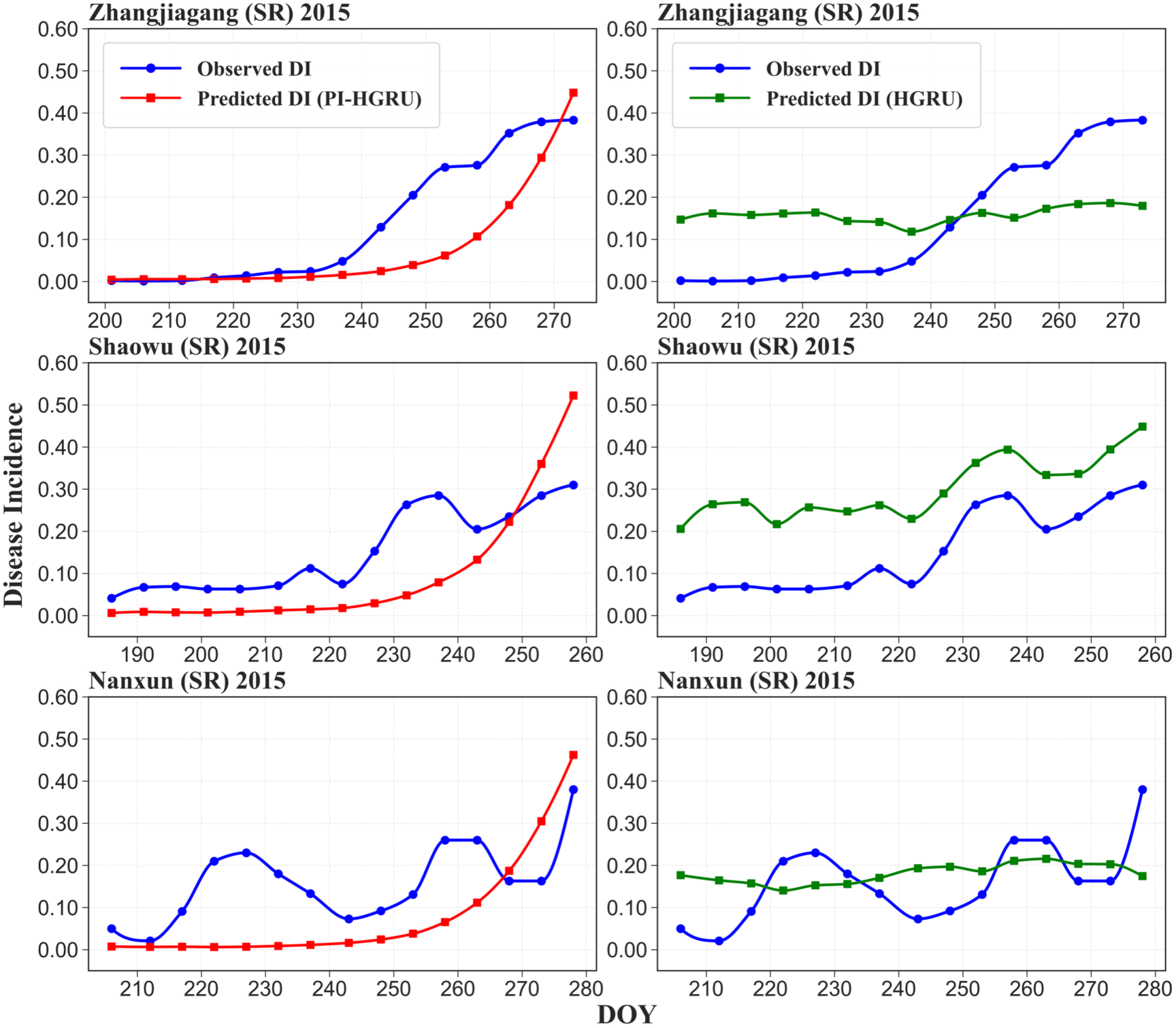
Observed and predicted rice sheath blight progression for representative growing-season sequences from the test set. The left panels show predictions from the physics-informed hybrid gated recurrent unit (PI-HGRU) model, while the right panels show predictions from the hybrid gated recurrent unit (HGRU) baseline. Each row represents the same growing-season sequence for the two models, allowing direct comparison of their temporal prediction patterns. Observed disease incidence is compared with the corresponding model predictions across the growing season. The figure illustrates differences in temporal prediction behavior between the two models under contrasting growing-season conditions. DI denotes disease incidence, DOY denotes day of year, and SR denotes single-season rice.

County-level prediction performance of PI-HGRU was further examined for early rice in 2015 and single-season rice in 2016 (Fig. 6). RMSE and MAE were relatively low across most evaluated counties in both test subsets (Fig. 6a–d). Higher errors occurred in a smaller number of counties and showed spatial variation across the study region. The spatial distributions of *r*² also varied among counties (Fig. 6e, f), with relatively high values in many evaluated counties and lower values in several localized areas. These results show that PI-HGRU maintained relatively consistent predictive performance across most evaluated counties, while local differences remained.

**Fig. 6.**
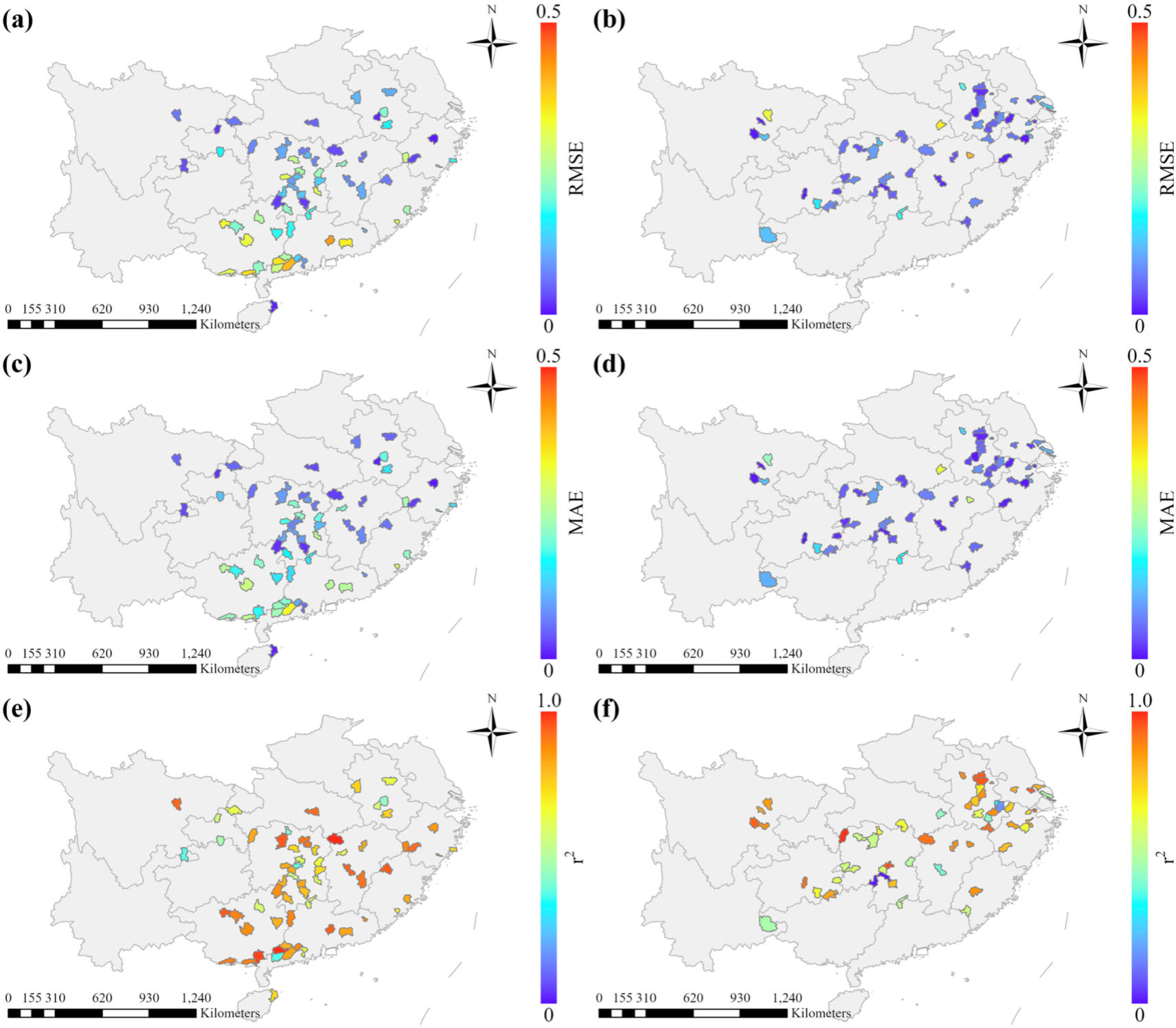
County-level prediction performance of the physics-informed hybrid gated recurrent unit. (PI-HGRU) model for early rice in 2015 and single-season rice in 2016 in China. Panels (a), (c), and (e) correspond to early rice in 2015, while panels (b), (d), and (f) correspond to single-season rice in 2016. Root mean square error (RMSE) is shown in panels (a) and (b), mean absolute error (MAE) in panels (c) and (d), and squared Pearson correlation coefficient (r²) in panels (e) and (f). Lower RMSE and MAE values and higher r² values indicate better county-level predictive performance.

Regional ShB risk predicted by PI-HGRU was visualized at four representative time points during the 2015 growing season, corresponding to day of year (DOY) 190, 210, 235, and 265 (Fig. 7). At DOY 190, all evaluated counties were classified as very low risk, with predicted disease incidence below 5% (Fig. 7a). By DOY 210, low-risk (5–10%) and moderate-risk (10– 20%) counties had emerged, indicating increasing spatial differentiation in predicted disease risk (Fig. 7b). At DOY 235, the risk distribution became more heterogeneous (Fig. 7c). Although most counties remained in the very low-risk category, moderate-risk, moderate-high-risk (20–40%), and high-risk (40–100%) counties occurred in several areas, including parts of the middle and lower reaches of the Yangtze River Basin. By DOY 265, fewer counties remained within their active growing seasons and were therefore included in the prediction map (Fig. 7d). Among the remaining counties, the proportions classified as moderate risk, moderate-high risk, or high risk increased, with several moderate-high- and high-risk counties occurring in the Yangtze River Delta and adjacent areas. Overall, the predicted risk maps showed a temporal transition from predominantly very low disease levels to increasingly heterogeneous risk distributions, accompanied by localized increases in predicted ShB incidence.

**Fig. 7.**
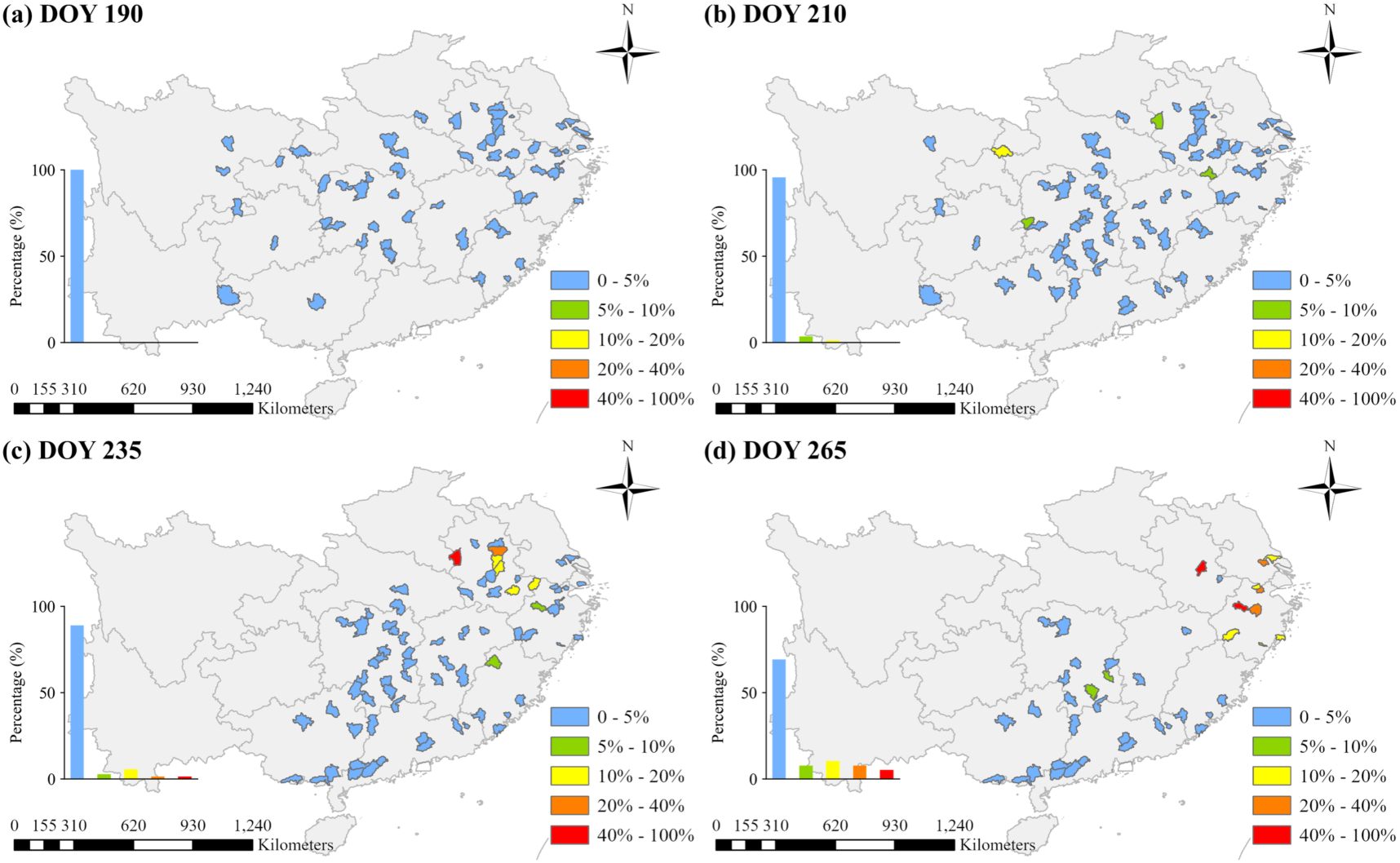
Predicted regional rice sheath blight risk at four representative time points during the 2015 growing season. Panels (a)–(d) show predictions at day of year (DOY) 190, 210, 235, and 265, respectively, illustrating changes in the spatial distribution of disease risk as the growing season progresses. Risk levels are classified according to predicted disease incidence as very low (0–5%), low (5–10%), moderate (10–20%), moderate-high (20–40%), and high (40–100%). The inset bar charts show the percentage of evaluated counties assigned to each risk class at the corresponding time point.

### 3.5. Model Interpretability and Environmental Response Analysis

Model interpretability analyses revealed clear differences in the contributions of environmental variables to ShB forecasting. PFI ranked VPD and land surface temperature (LST) as the two most influential predictors in the PI-HGRU model, followed by CL, precipitation, TP, TK, air temperature, soil water, and wind speed (Fig. S5). Remote sensing variables showed uneven contributions, with LST ranking highly, fraction of photosynthetically active radiation (FPAR) and leaf area index (LAI) showing moderate importance, and normalized difference water index (NDWI), enhanced vegetation index (EVI), and normalized difference vegetation index (NDVI) contributing comparatively little. Among the soil property variables, CL, TP, TK, available potassium (AK), pH, available phosphorus (AP), SOM, and available nitrogen (AN) all contributed to predictive performance.

The learned transmission rate *β*(*t*) showed distinct relationships with environmental statistics within the input time window (Fig. S6). Strong positive correlations were observed for LAI(max), Soil Water(max), Precip(min), LAI(std), Soil Water(mean), Soil Water(min), and LAI(mean), with *r* values ranging from 0.75 to 0.93. In contrast, wind-speed statistics showed consistently strong negative correlations with *β*(*t*), particularly Wind Speed(mean) (*r* = −0.95), Wind Speed(std) (*r* = −0.92), and Wind Speed(max) (*r* = −0.91). Several hydrothermal variables showed contrasting relationships among their temporal statistics. VPD(min) was strongly negatively correlated with *β*(*t*), whereas VPD(std) was positively correlated and VPD(max) showed little association. Minimum air temperature was positively correlated with *β*(*t*), whereas the mean, maximum, and standard deviation of air temperature were negatively correlated. Similarly, LST(mean), LST(max), and LST(min) were negatively correlated with *β*(*t*), while LST(std) showed a positive relationship.

## 4. Discussion

### 4.1. Environmental Drivers of ShB Forecasting

Protecting rice production from major diseases is key for global food security (Yu et al., 2026). Among these diseases, rice sheath blight (ShB) represents a tangible threat, for it is widespread and highly destructive, highlighting the need for reliable disease forecasting (Yellareddygari et al., 2014). Large-scale ShB forecasting, however, is complicated by substantial environmental heterogeneity. The development of this disease is strongly influenced by climatic and production conditions across growing environments (Savary et al., 2006). Thus, representing disease dynamics across such heterogeneous environments remains a major challenge in plant disease modelling (Cunniffe et al., 2015). To address this challenge, we developed the physics-informed hybrid gated recurrent unit (PI-HGRU) model by integrating multi-source environmental information with data-driven learning and a susceptible–exposed– infectious–removed (SEIR)-based epidemiological component. PI-HGRU improved forecasting performance relative to the purely data-driven baseline. The learned transmission rate also varied systematically with environmental conditions. These findings support the integration of epidemiological mechanisms with data-driven learning for disease forecasting (Ye et al., 2025).

Hydrothermal conditions were the dominant dynamic drivers in our ShB forecasting. We found that moisture conditions are important for ShB development and epidemic progression, in agreement with previous studies (Biswas et al., 2011). Indeed, rainfall can further promote the initiation and progression of this disease by maintaining favorable moisture conditions (Pal et al., 2017). Temperature also contributed substantially to our forecasting, although its relationships with the learned transmission rate varied among temporal statistics. In this context, field observations have shown that temperature can strongly influence the progression of ShB (Bhukal et al., 2015). More broadly, favorable temperature and moisture conditions jointly have been reported to contribute to environments becoming conducive to the development of ShB (Senapati et al., 2022).

The temporal statistics we used in our study suggest that preceding environmental conditions are key for the development of ShB. Indeed, we found that mean conditions, extremes, and short-term variability can provide complementary information beyond single-date observations. Previous studies have similarly linked preceding weather conditions with subsequent ShB development and forecasting (Biswas et al., 2011). Rainfall, temperature, and humidity have also been associated with the timing and progression of ShB epidemics (Pal et al., 2017). Similar use of preceding weather information has been reported in forecasting systems for other crop diseases (De Wolf et al., 2003). Together, these findings highlight the importance of incorporating historical environmental information to represent cumulative environmental effects and delayed disease responses during forecasting.

Canopy conditions and soil properties in our study provided additional environmental information for the forecasting of ShB. Their contributions may partly reflect their relationships with the canopy microenvironment in which disease develops. Dense rice canopies can increase humidity and leaf wetness, creating conditions favorable for fungal infection (Huber and Gillespie, 1992; Senapati et al., 2022). Soil nutrient conditions may further influence this microenvironment through their effects on crop growth and canopy structure. For example, nitrogen supply can alter canopy structure, tissue contact, and moisture retention, thereby affecting ShB spread (Savary et al., 1995). From the same perspective, the negative relationships between wind speed and the learned transmission rate may reflect increased canopy ventilation and reduced persistence of humid conditions. However, this interpretation requires further investigation because *β*(*t*), the time-varying transmission rate learned by the physics-informed component, is a model-derived quantity rather than a directly observed epidemiological parameter.

### 4.2. Performance and Interpretability of the PI-HGRU Model

PI-HGRU outperformed the purely data-driven hybrid gated recurrent unit (HGRU) baseline. This improvement highlights the potential value of integrating epidemiological mechanisms with data-driven learning for disease forecasting. Mechanistic models explicitly represent disease transmission processes, but their performance can depend on model structure, parameterization, and data availability (Cunniffe et al., 2015). Data-driven models can capture complex nonlinear relationships from observations, but generally lack explicit representations of underlying epidemic processes (Ye et al., 2025). In contrast, physics-informed modelling provides a way to integrate these complementary sources of information (Qian et al., 2025). In PI-HGRU, the SEIR-based component represents disease progression, while the data-driven branch captures complex environmental relationships. Their contributions are adaptively combined through the dynamic fusion mechanism. We argue that this integration likely contributed to the improved forecasting performance across heterogeneous growing-season sequences.

The physics-informed component of our model provides key information for interpreting our predictions. Previous studies have shown that epidemiological mechanisms can be incorporated into data-driven models while allowing transmission parameters to be learned from observations (Ye et al., 2025). Environmental covariates can also be incorporated into physics-informed neural networks to represent variation in disease transmission (Qian et al., 2025). In our PI-HGRU model, the learned transmission rate *β*(*t*) provides an intermediate representation of how disease transmission varies with environmental conditions. Analysis of *β*(*t*) therefore complements permutation feature importance (PFI), which quantifies the contribution of individual environmental variables to predictive performance. However, *β*(*t*) is a model-derived quantity, rather than a directly observed epidemiological parameter. As such, relationships between *β*(*t*) and environmental variables should be interpreted as model-based associations rather than causal effects, particularly given uncertainties in data, parameterization, and model structure (D’Agostino McGowan et al., 2021).

### 4.3. Application Potential, Limitations, and Future Directions

The regional forecasting capability of PI-HGRU highlights its potential for large-scale ShB risk monitoring and decision support. Spatially continuous forecasts can complement field disease surveys by identifying areas with increasing disease risk and prioritizing locations for further inspection. Regional disease assessments based on remote sensing and meteorological information have shown potential for supporting ShB monitoring and control (Zhang et al., 2023). Weather-based ShB forecasting has also been applied to provide early warning and improve disease control (Chu et al., 2023). More broadly, regional crop disease forecasts can provide risk alerts for agricultural decision support (Skelsey, 2021). Because the temporal inputs required by PI-HGRU can be derived from routinely available meteorological and remote sensing data, the framework provides a practical basis for updating regional ShB risk information throughout the growing season.

Several limitations should be considered when interpreting our model predictions. In our study, disease observations were collected at discrete locations and dates, whereas environmental variables were aggregated to county-scale conditions. This approach could have obscured local variation in crop and disease conditions. Important management factors affecting the development of ShB, including cultivar susceptibility, planting density, fertilization, and fungicide application (Tang et al., 2007; Wu et al., 2013), were also unavailable across regions and years. In addition, the SEIR-based component simplifies ShB epidemic development by representing environmental variation primarily through the learned transmission rate *β*(*t*). Such simplification is inherent in epidemiological models that represent selected processes of complex biological systems (Cunniffe et al., 2015; Savary et al., 2012). Future work should evaluate the framework in additional regions and years, incorporate higher-resolution crop and management information, and investigate whether additional epidemiological parameters can be dynamically estimated or constrained using independent biological knowledge.

## 5. Conclusion

This study addresses the challenge of large-scale spatiotemporal forecasting of rice sheath blight (ShB) under complex environmental conditions. To do so, we developed a physics-informed hybrid gated recurrent unit (PI-HGRU) model that integrates epidemiological mechanisms with deep learning-based spatiotemporal modelling. The time-window sensitivity analysis identified a 45-day input time window with a 7-day lag time as the optimal forecasting configuration, indicating that both cumulative environmental effects and delayed disease responses should be considered in regional disease forecasting. Under this configuration, the proposed model achieved a satisfactory predictive performance across the major rice-growing regions of southern China. Compared with the hybrid gated recurrent unit (HGRU) baseline model, PI-HGRU improved the squared Pearson correlation coefficient (*r*²) by 22.8% while reducing root mean square error (RMSE) and mean absolute error (MAE) by 10.2% and 18.0%, respectively, demonstrating that the incorporation of epidemiological constraints effectively enhanced forecasting performance across different regions and years. In addition, our model outputs effectively captured the temporal progression and spatial differentiation of disease development at the regional scale. The learned transmission rate *β*(*t*) further enabled quantitative analysis of the relationships between disease transmission and environmental drivers, thereby improving model interpretability. Overall, our study demonstrates the feasibility of integrating epidemiological mechanisms with deep learning for large-scale spatiotemporal disease forecasting. The proposed framework provides an effective methodological reference for developing interpretable and robust disease forecasting models and offers a promising foundation for regional disease forecasting, risk warning, and precision disease management.

## Supporting information

SOMs

## Acknowledgements

This work was supported by Zhejiang Provincial Natural Science Foundation of China (Grant no. LR25D010003); National Natural Science Foundation of China (Grant no. 42401400); China Scholarship Council (Grant no. 202608330171).

