## Supplementary material for "A physics-informed hybrid deep learning model for spatiotemporal rice disease prediction using multi-source data": SOMs

**Table of references** **Page**

1. Correlations among meteorological statistical features 2

2. Correlations among soil property variables 4

3. Permutation feature importance for input feature selection 5

4. Sequence-level model performance distributions 7

5. Permutation feature importance of environmental variables in PI-HGRU 8

6. Environmental relationships of the learned transmission rate $\beta(t)$ 10

### 1. Correlations among meteorological statistical features

**Fig. S1. Pearson correlation matrix of growing-season meteorological statistical features used for feature selection.** Pairwise Pearson correlation coefficients quantify the strength and direction of linear relationships among statistical features derived from the meteorological variables. The color scale represents correlation coefficients ranging from negative to positive relationships, with larger absolute values indicating stronger correlations. The matrix is used to identify highly correlated features and reduce redundancy before subsequent feature importance analysis. **Abbreviations:** Temp, 2-m air temperature; Precip, total daily precipitation; Dewpoint, 2-m dewpoint temperature; Soil Temp, soil temperature (0–7 cm); Soil Water, volumetric soil water (0–7 cm); Wind Speed, 10-m wind speed; SSRD, surface solar radiation downwards; RH, relative humidity; VPD, vapor pressure deficit. The suffixes mean, std, median, min, max, and cv denote mean, standard deviation, median, minimum, maximum, and coefficient of variation, respectively.

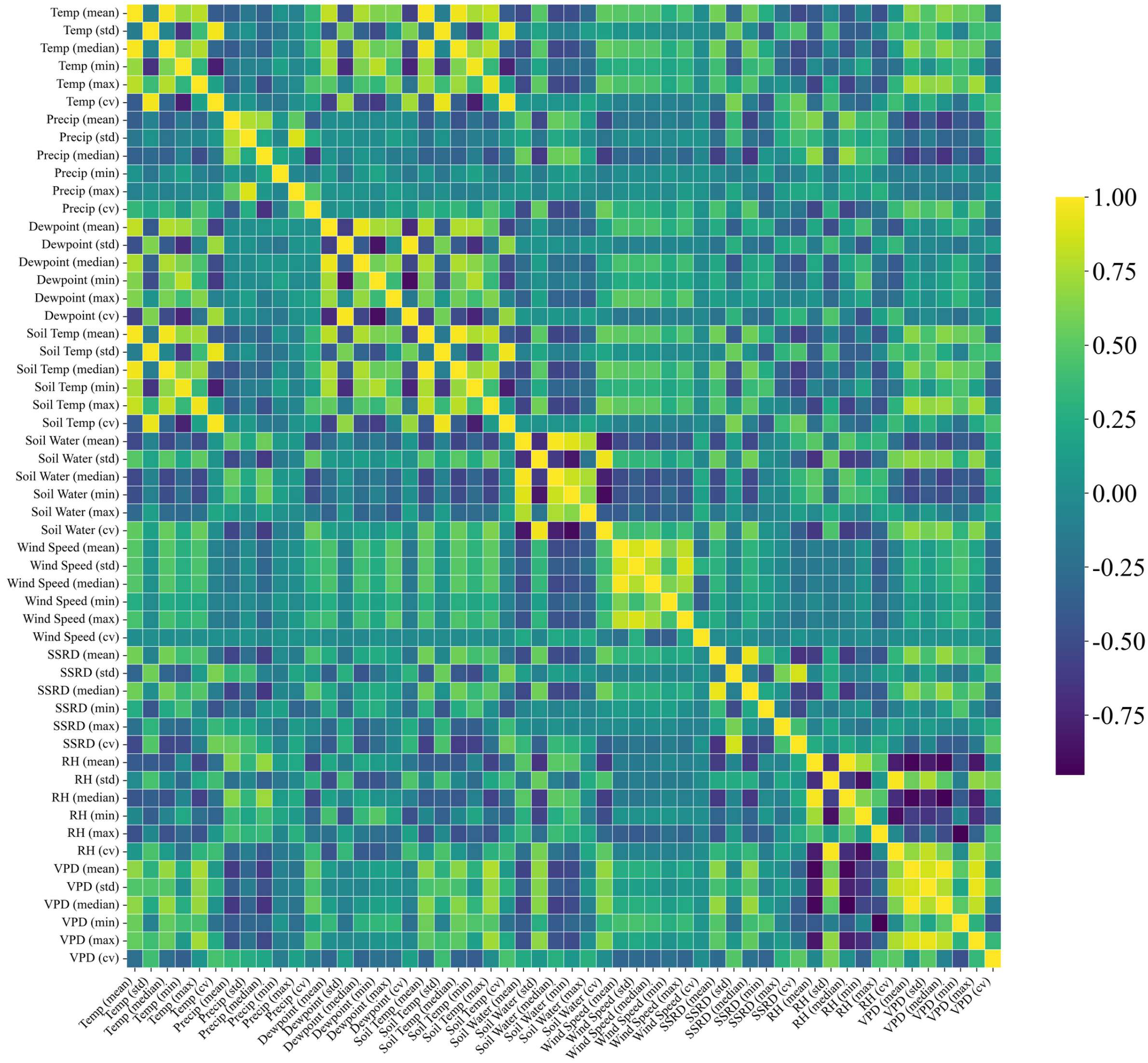
Temp (mean)
Temp (std)
Temp (median)
Temp (min)
Temp (max)
Temp (cv)
Precip (mean)
Precip (std)
Precip (median)
Precip (min)
Precip (max)
Precip (cv)
Dewpoint (mean)
Dewpoint (std)
Dewpoint (median)
Dewpoint (min)
Dewpoint (max)
Dewpoint (cv)
Soil Temp (mean)
Soil Temp (std)
Soil Temp (median)
Soil Temp (min)
Soil Temp (max)
Soil Temp (cv)
Soil Water (mean)
Soil Water (std)
Soil Water (median)
Soil Water (min)
Soil Water (max)
Soil Water (cv)
Wind Speed (mean)
Wind Speed (std)
Wind Speed (median)
Wind Speed (min)
Wind Speed (max)
Wind Speed (cv)
SSRD (mean)
SSRD (std)
SSRD (median)
SSRD (min)
SSRD (max)
SSRD (cv)
RH (mean)
RH (std)
RH (median)
RH (min)
RH (max)
RH (cv)
VPD (mean)
VPD (std)
VPD (median)
VPD (min)
VPD (max)
VPD (cv)
1.00
0.75
0.50
0.25
0.00
-0.25
-0.50
-0.75

### 2. Correlations among soil property variables

**Fig. S2. Pearson correlation matrix of soil property variables used for feature selection.** Pairwise Pearson correlation coefficients quantify the strength and direction of linear relationships among the soil property variables. The color scale represents correlation coefficients ranging from negative to positive relationships, with larger absolute values indicating stronger correlations. The matrix is used to identify highly correlated soil property variables and reduce redundancy before subsequent feature importance analysis. **Abbreviations:** AK, available potassium; AN, available nitrogen; AP, available phosphorus; CL, clay content; pH, soil pH; SOM, soil organic matter; TK, total potassium; TN, total nitrogen; TP, total phosphorus.

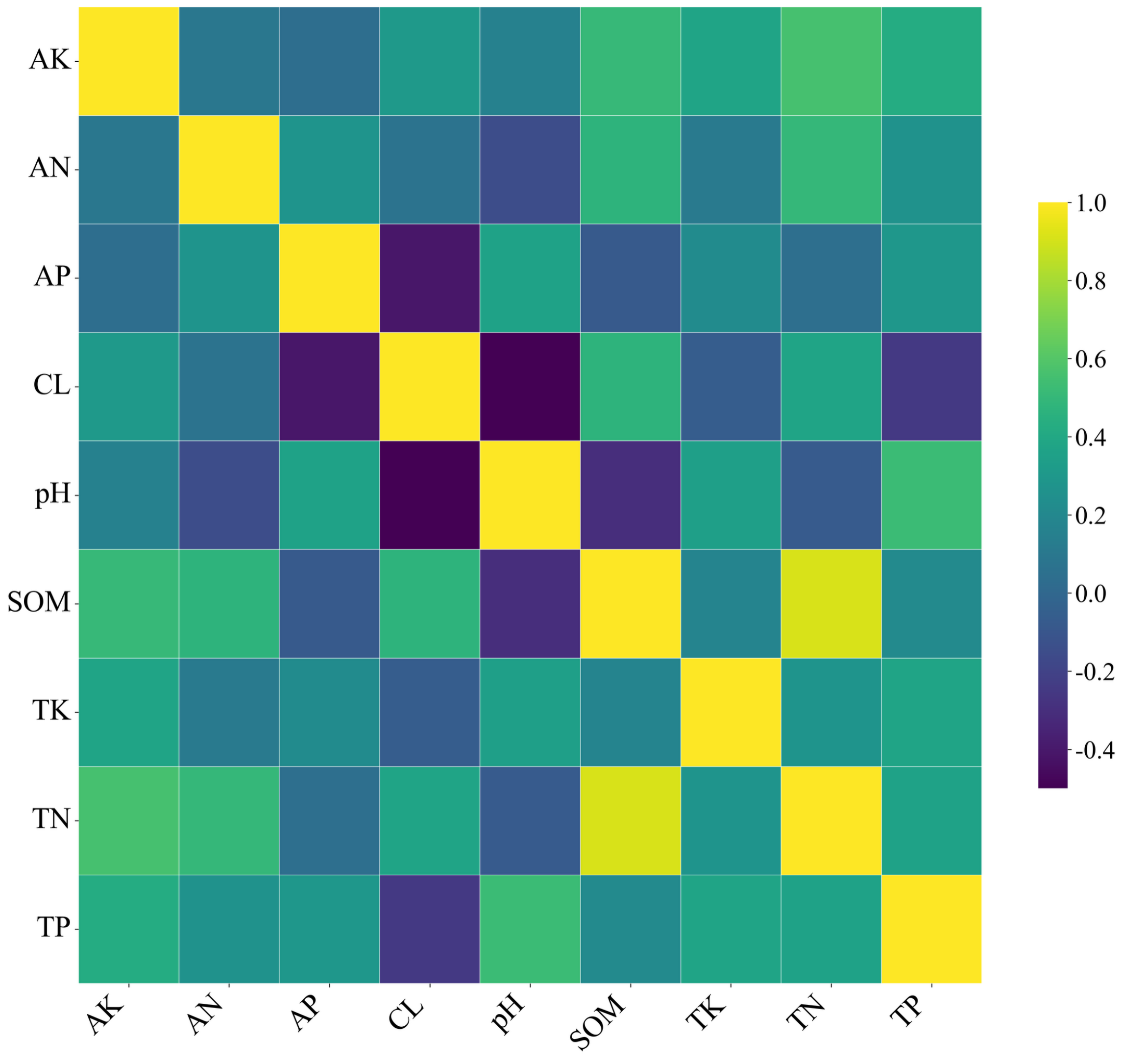

### 3. Permutation feature importance for input feature selection

**Fig. S3. Permutation feature importance (PFI) ranking of candidate features used for rice sheath blight forecasting.** Feature importance is quantified by the change in prediction error after individual features are permuted, with larger positive scores indicating greater contributions to model performance. Colors distinguish meteorological statistical features, soil property features, and principal components derived from remote sensing features. The left panel shows features within the top 50% of the PFI ranking, while the right panel shows the remaining features. Features in the top 50% are used to determine the variables retained for subsequent model construction. **Abbreviations:** VPD, vapor pressure deficit; TK, total potassium; SSRD, surface solar radiation downwards; TP, total phosphorus; CL, clay content; AK, available potassium; RS_PC, principal component derived from remote sensing statistical features; AN, available nitrogen; AP, available phosphorus; pH, soil pH; SOM, soil organic matter; Dewpoint, 2-m dewpoint temperature; Wind Speed, 10-m wind speed; Soil Water, volumetric soil water (0–7 cm); Precip, total daily precipitation; Temp, 2-m air temperature. The suffixes mean, std, min, and max denote mean, standard deviation, minimum, and maximum, respectively.

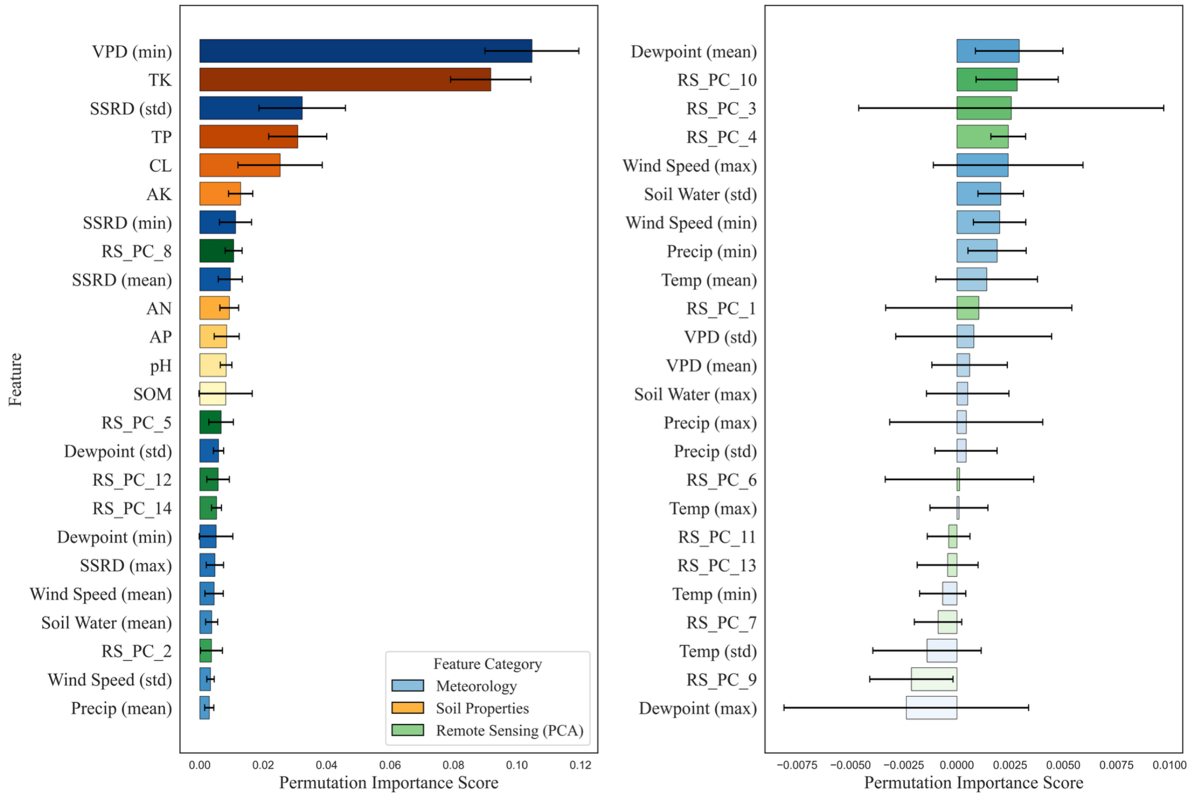
Feature
VPD (min)
TK
SSRD (std)
TP
CL
AK
SSRD (min)
RS_PC_8
SSRD (mean)
AN
AP
pH
SOM
RS_PC_5
Dewpoint (std)
RS_PC_12
RS_PC_14
Dewpoint (min)
SSRD (max)
Wind Speed (mean)
Soil Water (mean)
RS_PC_2
Wind Speed (std)
Precip (mean)
Feature Category
Meteorology
Soil Properties
Remote Sensing (PCA)
0.00 0.02 0.04 0.06 0.08 0.10 0.12
Permutation Importance Score
Dewpoint (mean)
RS_PC_10
RS_PC_3
RS_PC_4
Wind Speed (max)
Soil Water (std)
Wind Speed (min)
Precip (min)
Temp (mean)
RS_PC_1
VPD (std)
VPD (mean)
Soil Water (max)
Precip (max)
Precip (std)
RS_PC_6
Temp (max)
RS_PC_11
RS_PC_13
Temp (min)
RS_PC_7
Temp (std)
RS_PC_9
Dewpoint (max)
−0.0075 −0.0050 −0.0025 0.0000 0.0025 0.0050 0.0075 0.0100
Permutation Importance Score

### 4. Sequence-level model performance distributions

**Fig. S4. Distributions of sequence-level prediction performance for the physics-informed hybrid gated recurrent unit (PI-HGRU) model and the hybrid gated recurrent unit (HGRU) baseline on the 2015–2016 test set.** Root mean square error (RMSE), mean absolute error (MAE), and squared Pearson correlation coefficient ($r^2$) are calculated separately for each growing-season sequence. Sequence-level metrics from five independent runs are pooled to characterize the distribution of model performance. Lower RMSE and MAE values and higher $r^2$ values indicate better predictive performance.

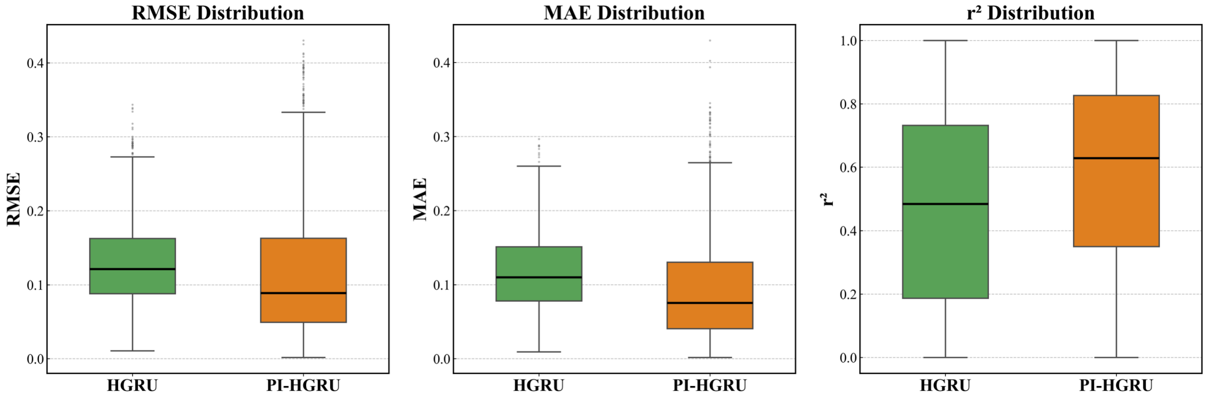

### 5. Permutation feature importance of environmental variables in PI-HGRU

**Fig. S5. Permutation feature importance (PFI) of environmental variables in the physics-informed hybrid gated recurrent unit (PI-HGRU) model.** Feature importance is quantified by the increase in root mean square error after individual environmental variables are permuted, with larger positive values indicating greater contributions to model performance. Colors distinguish meteorological, remote sensing, and soil property variables. **Abbreviations:** VPD, vapor pressure deficit; LST, land surface temperature (day-time); CL, clay content; Precip, total daily precipitation; TP, total phosphorus; TK, total potassium; Temp, 2-m air temperature; Soil Water, volumetric soil water (0–7 cm); Wind Speed, 10-m wind speed; AK, available potassium; Dewpoint, 2-m dewpoint temperature; pH, soil pH; AP, available phosphorus; SOM, soil organic matter; AN, available nitrogen; FPAR, fraction of photosynthetically active radiation; LAI, leaf area index; SSRD, surface solar radiation downwards; NDWI, normalized difference water index; EVI, enhanced vegetation index; NDVI, normalized difference vegetation index.

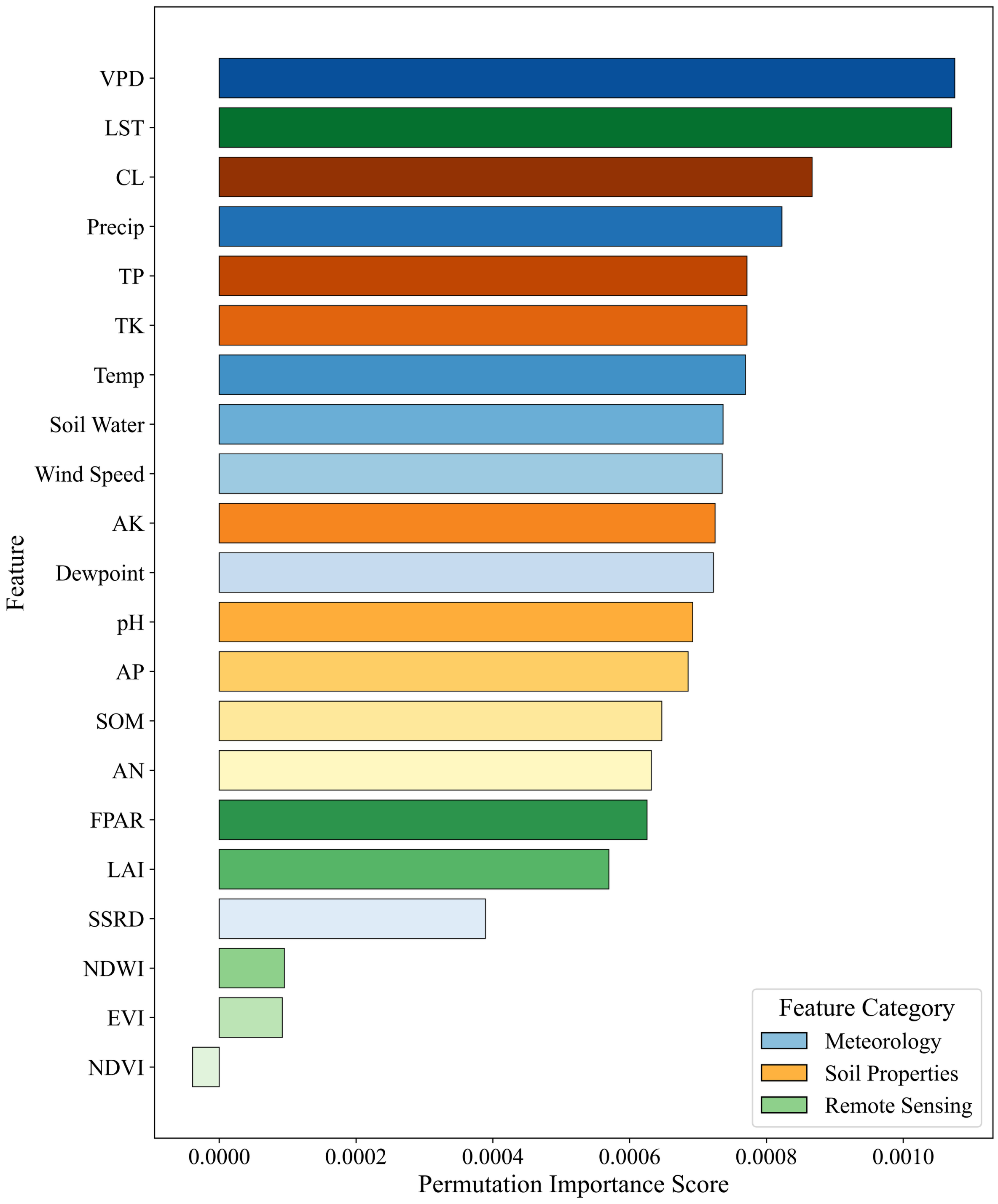
VPD
LST
CL
Precip
TP
TK
Temp
Soil Water
Wind Speed
AK
Dewpoint
pH
AP
SOM
AN
FPAR
LAI
SSRD
NDWI
EVI
NDVI
Feature
Feature Category
Meteorology
Soil Properties
Remote Sensing
0.0000
0.0002
0.0004
0.0006
0.0008
0.0010
Permutation Importance Score

### 6. Environmental relationships of the learned transmission rate $\boldsymbol{\beta(t)}$

**Fig. S6. Pearson correlations between the learned transmission rate $\boldsymbol{\beta(t)}$ and environmental statistics within the historical input window.** For each temporal environmental variable, the mean, standard deviation, minimum, and maximum are calculated over the corresponding input window. Pearson correlation coefficients quantify the linear associations between these historical environmental statistics and the corresponding $\beta(t)$ values. Positive and negative coefficients indicate positive and negative associations, respectively, while larger absolute values indicate stronger linear relationships. **Abbreviations:** LAI, leaf area index; Soil Water, volumetric soil water (0–7 cm); Precip, total daily precipitation; FPAR, fraction of photosynthetically active radiation; EVI, enhanced vegetation index; LST, land surface temperature (day-time); NDVI, normalized difference vegetation index; Dewpoint, 2-m dewpoint temperature; VPD, vapor pressure deficit; Temp, 2-m air temperature; NDWI, normalized difference water index; SSRD, surface solar radiation downwards; Wind Speed, 10-m wind speed. The suffixes mean, std, min, and max denote mean, standard deviation, minimum, and maximum, respectively.

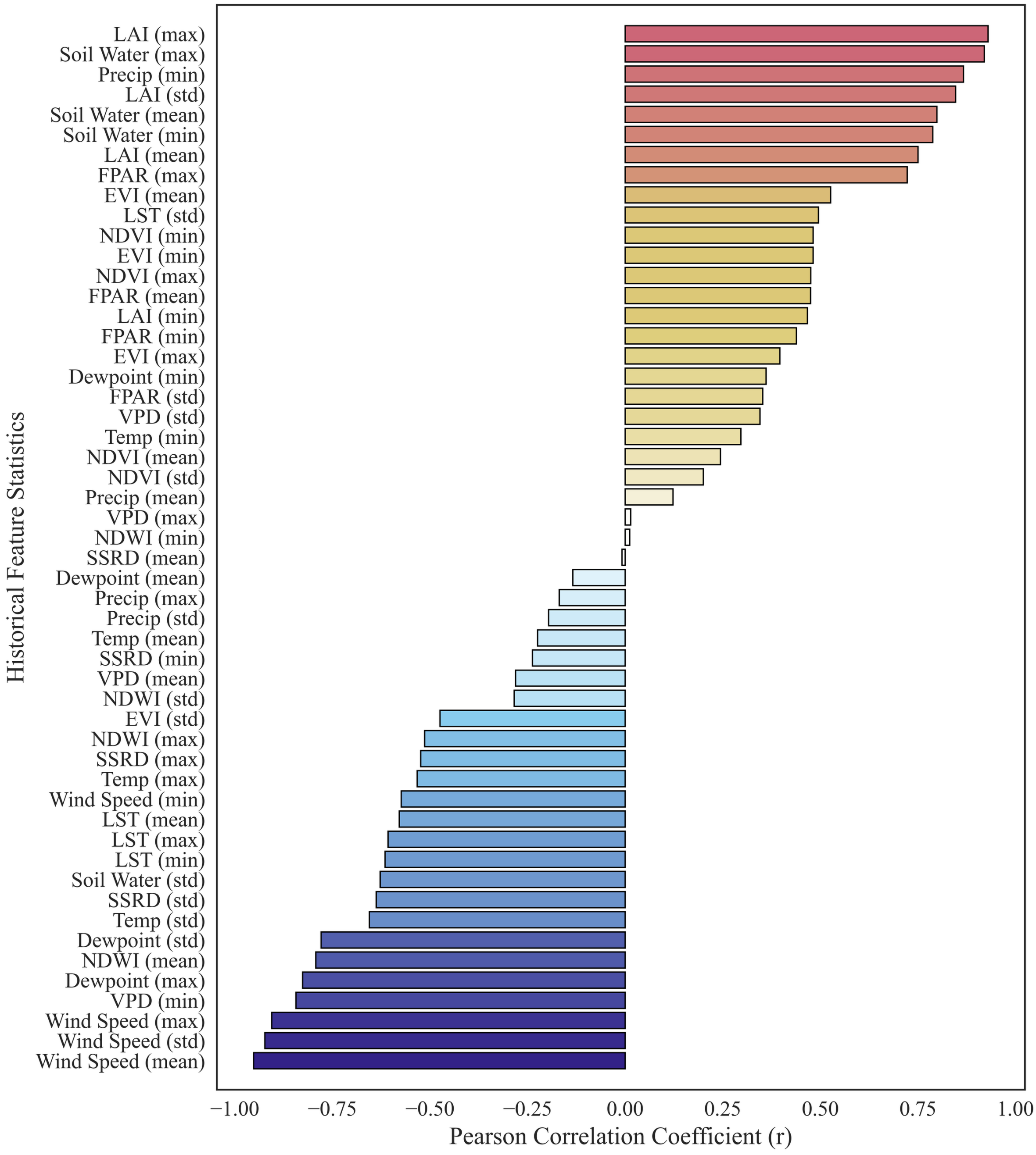
LAI (max)
Soil Water (max)
Precip (min)
LAI (std)
Soil Water (mean)
Soil Water (min)
LAI (mean)
FPAR (max)
EVI (mean)
LST (std)
NDVI (min)
EVI (min)
NDVI (max)
FPAR (mean)
LAI (min)
FPAR (min)
EVI (max)
Dewpoint (min)
FPAR (std)
VPD (std)
Temp (min)
NDVI (mean)
NDVI (std)
Precip (mean)
VPD (max)
NDWI (min)
SSRD (mean)
Dewpoint (mean)
Precip (max)
Precip (std)
Temp (mean)
SSRD (min)
VPD (mean)
NDWI (std)
EVI (std)
NDWI (max)
SSRD (max)
Temp (max)
Wind Speed (min)
LST (mean)
LST (max)
LST (min)
Soil Water (std)
SSRD (std)
Temp (std)
Dewpoint (std)
NDWI (mean)
Dewpoint (max)
VPD (min)
Wind Speed (max)
Wind Speed (std)
Wind Speed (mean)
Historical Feature Statistics
−1.00
−0.75
−0.50
−0.25
0.00
0.25
0.50
0.75
1.00
Pearson Correlation Coefficient (r)
